# Cutaneous inflammation accelerates the premalignant expansion of melanocytes bearing oncogenic mutations

**DOI:** 10.64898/2026.05.27.728115

**Authors:** Diana Tran, Anastasiia Vaska, Tyler El Rayes, Natalie Lovinger, Yassmin A. Elbanna, Esther Lee, Christin E. Burd, Jonathan H. Zippin, Morgan Huse

## Abstract

How the cutaneous microenvironment influences early melanomagenesis is poorly understood. Here, we assessed the effects of three immune perturbations on premalignant melanocyte expansion in an autochthonous mouse model of disease. Depletion of regulatory T (T_reg_) cells markedly accelerated melanoproliferation, an unexpected phenotype that was associated with monocyte and macrophage infiltration, the production of inflammatory and angiogenic factors, and vascular leakage. In line with these observations, single cell transcriptomic analysis of T_reg_ cell deficient skin revealed robust accumulation of monocyte-derived macrophages with tissue remodeling characteristics. Acute UV irradiation and 2,4-dinitrofluorobenzene (DNFB)-induced contact hypersensitivity had analogous effects on both the cellular microenvironment of the skin and the expansion of local premalignant melanocytes. Treatment with the anti-inflammatory agent dexamethasone attenuated DNFB-induced melanocyte expansion and vascular remodeling. Collectively, these results identify a conserved inflammatory axis linked to the early outgrowth of oncogenic melanocytes in the skin.

## Introduction

Melanoma is the most lethal form of skin cancer, accounting for approximately 75% of skin cancer–related deaths^1,2^. It develops through the malignant transformation of melanocytes, which proliferate uncontrollably and acquire metastatic potential^3^. Whereas numerous targeted small molecule inhibitors and immune-activating therapeutics are FDA approved to treat metastatic disease, only 50% of patients achieve durable response^4^. Thus, it is critical to better understand the development of melanoma to identify improved methods for diagnosis and treatment. Among the major drivers of melanomagenesis are activating somatic mutations in *NRAS* and *BRAF*, which are often coupled with loss-of-function mutations in tumor suppressors like *TP53* and *PTEN*^5,6^. However, these same genetic alterations are found in benign melanocytic nevi^7–10^, implying that factors other than oncogenic melanocyte mutations are also necessary for malignant progression.

The immune system represents an obvious source for melanocyte-extrinsic determinants of melanoma. Multiple immune cell types, including natural killer cells and CD8^+^ cytotoxic T lymphocytes, destroy cancer cells directly, and this anti-tumor activity is widely thought to eliminate many nascent tumors before they become clinically detectable^11,12^. Anti-tumor immunity is particularly relevant for melanoma, which can be successfully treated with immune checkpoint blockade (ICB) antibodies that act by derepressing existing clones of tumor-specific T cells^13,14^. Accordingly, the capacity of mutant melanocytes to overcome or evade anti-tumor lymphocytes would presumably be crucial for their outgrowth in the skin.

Regulatory T (T_reg_) cells, which are defined by expression of the transcription factor Foxp3, function as critical suppressors of lymphocyte activation in both homeostatic and disease contexts^15^. As such, they represent an intriguing candidate immune evasion mechanism for melanoma. Indeed, multiple lines of evidence link T_reg_ cell activity to melanoma outgrowth. Intratumoral T_reg_ cell accumulation has been associated with disease progression in the clinic, and T_reg_ cell depletion inhibits melanoma outgrowth in transplantable models of the disease^16–19^. However, the role of T_reg_ cells during the early, premalignant stage of melanocyte expansion has not been examined.

Beyond anti-tumor immunity, the immune system can also influence tumor progression via tissue inflammation^12,20^. Inflammatory responses are typically triggered by the release of activating cytokines and chemokines in peripheral tissues^21,22^. This drives the recruitment of a variety of innate immune cells from the blood, among them neutrophils and Ly6C^+^ monocytes, the latter of which rapidly differentiate into inflammatory macrophages. Infiltrating myeloid cells further amplify the inflammatory milieu and release factors that promote tissue remodeling. A key component of this remodeling response is angiogenesis, a process in which initial vascular destabilization is followed by the sprouting of new vessels^23^. Importantly, myeloid-driven inflammation and angiogenesis have both been shown to support tumor progression by providing the space and nutrients necessary for cancer cell growth^20,23^.

The link between inflammation and cancer may be particularly relevant to melanoma because ultraviolet (UV) light, the primary environmental driver of this disease, is known to elicit robust cutaneous inflammation and vascular remodeling^24–26^. UV is canonically thought to drive melanomagenesis by generating oncogenic mutations in melanocyte DNA^6,27–30^. While UV-induced mutations clearly contribute to disease progression, emerging evidence indicates that UV may also promote disease via non-mutagenic pathways^31–35^. Mouse models of melanoma have shown that a single dose of UV is sufficient to induce melanomagenesis with no significant increase in oncogenic UV signature mutations^36,37^. Furthermore, acute high doses of UV (e.g., blistering sunburn) are a more significant risk factor for melanoma than chronic UV exposure^31,38–41^, despite presumably generating less oncogenic DNA damage. In light of these data, it is tempting to speculate that inflammatory tissue remodeling induced by UV light might create local conditions that are conducive to melanomagenesis. Intriguingly, UV-induced inflammation is also associated with cutaneous T_reg_ cell expansion and the suppression of adaptive immunity in the skin^42–44^. Hence, UV could support melanoma outgrowth by driving both tissue remodeling and immune suppression.

To study the interplay between incipient melanoma and cutaneous inflammation, we subjected an autochthonous murine model of melanoma (*LSL-Braf^V600E^*;*Pten^fl/fl^;Tyr::CreERT2* mice) to three distinct inflammatory immune perturbations: transient T_reg_ cell depletion, acute UV-B irradiation, and 2,4-dinitrofluorobenzene (DNFB)-induced contact hypersensitivity. Each of these perturbations accelerated premalignant melanocyte outgrowth, which was unexpected given that both T_reg_ cell depletion and DNFB treatment markedly enhanced conventional T (T_conv_) cell infiltration into the skin. Detailed analysis of each inflammatory response revealed a shared cellular and molecular signature comprising myeloid infiltration, characteristic cytokines and tissue remodeling factors, and vascular permeability. Importantly, pharmacological suppression of this inflammatory signature mitigated melanocyte expansion, indicating that it plays a central role in early disease progression. These results are consistent with a model in which oncogenic mutation synergizes with tissue destabilization to drive melanomagenesis.

## Results

### Accumulation of T_reg_ cells in early melanoproliferative lesions

To explore early interactions between oncogene-containing melanocytes and the cutaneous immune system, we employed an established autochthonous model of melanoma: *LSL-Braf^V600E^*;*Pten^fl/fl^;Tyr::CreERT2* (*BPT)* mice^45^. Topical treatment of these animals with 4-hydroxytamoxifen (4-HT) induces the expression of oncogenic Braf^V600E^ concomitantly with the deletion of the tumor suppressor Pten in melanocytes, which proceed over the ensuing ∼8 weeks to form melanoma. We focused on the glabrous skin of the mouse ear because it is conducive to intravital microscopy and because it closely mimics the architecture of human skin.

Our initial studies utilized *BPT* mice bearing an *LSL-TdTomato* allele, which generate TdTomato^+^ melanocytes after 4-HT treatment. This labelling strategy was selected to facilitate the imaging of melanocytes *in situ* and to enable detection of melanocytic material uptake by immune cells^46^. *BPT;LSL-TdTomato* mice, along with *Tyr::CreER;LSL-TdTomato* controls lacking tumorigenic alleles, were subjected to 4-HT ear painting, and immune cells in the ear skin and the draining lymph nodes (dLN) were analyzed by flow cytometry 35 days later (Fig. 1A, Fig. 1 - supplement 1A-B). At this time point, *BPT* mice exhibited substantial skin darkening due to melanocyte outgrowth, but they had not yet developed obvious tumors (Fig. 1A). Multiple CD45^+^ immune cell types were enriched in *BPT* ears relative to wild type controls, including T cells (CD3^+^), monocytes (CD11b^+^Ly6G^-^F4/80^+^MHCII^-^CCR2^+^), and macrophages (CD11b^+^Ly6G^-^F4/80^+^) (Fig. 1B). Conventional type I (CD11c^+^MHCII^+^F4/80^-^CD103^+^CD11b^-^) and type II (CD11c^+^MHCII^+^F4/80^-^CD103^-^CD11b^+^) dendritic cells (cDC1 and cDC2) were also more abundant in BPT skin (Fig. 1C). Infiltrating T cells were predominantly CD4^+^, with only small numbers of CD8^+^ T cells detected (Fig. 1B). Notably, approximately one quarter of CD4^+^ T cells in BPT skin expressed Foxp3, indicative of a substantive T_reg_ cell response (Fig. 1B). Increased T_conv_ and T_reg_ cell numbers were also observed in the *BPT* dLN, but these changes were more modest and fell short of statistical significance (Fig. 1 - supplement 1C-D).

**Figure 1.**
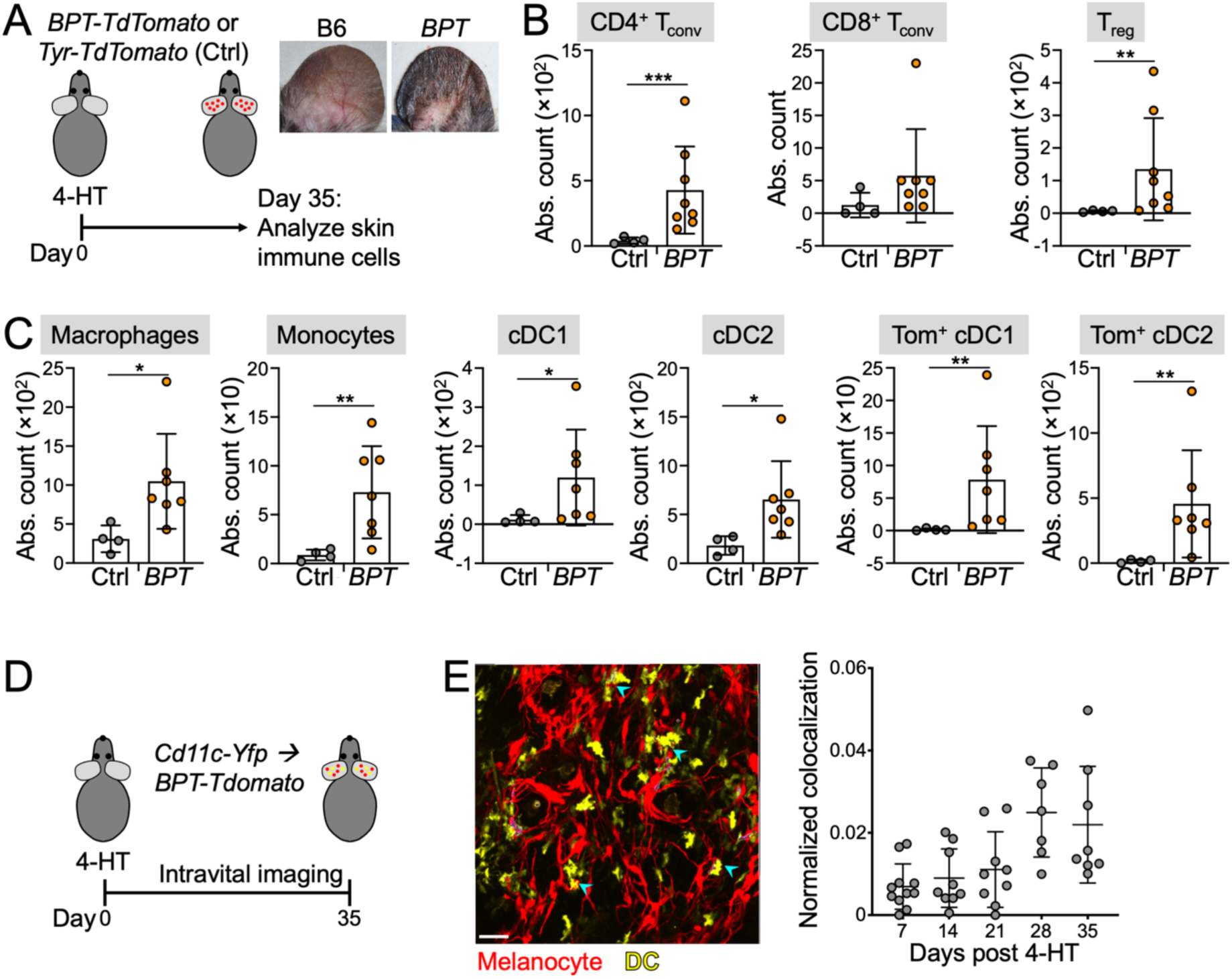
Premalignant melanocyte outgrowth is accompanied by cutaneous immune infiltration. (A-C) *BPT-TdTomato* and *Tyr-TdTomato* control mice were 4-HT-painted and CD45^+^ cells in the skin enumerated after 35 days by flow cytometry. (A) Schematic diagram of the experimental approach. Insets show representative melanoproliferation in *BPT* ears 1 month after 4-HT painting. (B) Quantification of the indicated T cell subsets. (C) Quantification of the indicated myeloid cell subsets. Tom^+^ = TdTomato positive. In B and C, *, **, and *** denote P ≤ 0.05, P < 0.01, and P < 0.001, respectively, calculated by lognormal Welch’s t-test, except in the case of Tom^+^ cDC1 and Tom^+^ cDC2, where Mann-Whitney test was applied. (D-E) *BPT-TdTomato* mice reconstituted with *Cd11c-yfp* bone marrow were 4-HT-painted and subjected to weekly two-photon imaging for 35 days. (D) Schematic diagram of the experimental approach. (E) Representative image of mutant melanocytes (red) and DCs (yellow) in the skin, with DC-melanocyte interactions indicated by cyan arrowheads. Scale bar = 50 μm. (F) Quantification of DC-melanocyte interaction frequency, normalized to the total number of DCs in each image. Error bars denote SD.

To assess the antigen presentation pathway underlying the enhanced T cell infiltration observed in *BPT* skin, we quantified TdTomato fluorescence in patrolling DC subsets. Little to no TdTomato uptake was observed in control mice, despite the fact that the unmutated melanocytes in these animals were TdTomato^+^. By contrast, ∼50% of cutaneous cDC1s and ∼80% of cDC2s in *BPT* ears contained melanocyte material (Fig. 1C). A substantial number of TdTomato^+^ cDC1s and cDC2s were also observed in the dLNs of 4-HT-treated *BPT* mice (Fig. 1 - supplement 1D), implying that both subsets were potentially capable of presenting mutant melanocyte antigen to naive T cells. To explore a potential basis for melanocyte sampling by DCs, we reconstituted *BPT*;*LSL-TdTomato* mice with *CD11c-YFP* bone marrow, enabling simultaneous two-photon imaging of YFP^+^ DCs and TdTomato^+^ melanocytes in premalignant ears (Fig. 1D). In these experiments, DCs initiated contact with melanocytes within three days of 4-HT treatment, and their interaction frequency trended upward over time as the melanocytes expanded (Fig. 1E). Hence, cutaneous DCs take up material from mutant melanocytes and traffic it to the dLN. Taken together with the increased accumulation of cutaneous T_conv_ cells in premalignant BPT skin, these results suggest that melanocyte specific T_conv_ and T_reg_ cells may be primed in the early stages of mutant melanocyte outgrowth.

### T_reg_ cells restrain oncogenic melanocyte outgrowth in the skin

The influx of both T_conv_ and T_reg_ cells into *BPT* skin led us to hypothesize that anti-melanocyte responses mediated by the former might be suppressed by the latter. To test this hypothesis, we prepared *BPT* mice containing the *Foxp3-DTR* allele, which drives diphtheria toxin (DT) receptor expression in T_reg_ cells and thereby sensitizes them to transient depletion by DT^47^. In our experiments, T_reg_ cells were depleted with three injections of DT delivered at 48-hour intervals. This treatment scheme reduced circulating T_reg_ cells to background levels for at least 10 days (Fig. 2 - supplement 1A-B). Cohorts of *BPT;Foxp3-DTR* animals were subjected to three different T_reg_ cell depletion regimens, starting six days before (early), two days after (intermediate), or eight days after (late) 4-HT painting (Fig. 2A). Surprisingly, none of these regimens inhibited 4-HT-induced melanoproliferation. Instead, we found that T_reg_ cell depletion, and in particular early T_reg_ cell depletion, enhanced the outgrowth response, which we visualized photographically and quantified by qRT-PCR of the melanocyte-specific marker *Tyrp1* (Fig. 2B-C). These results indicate that T_reg_ cells constrain premalignant melanocyte expansion during a temporal window closely aligned with the induction of tumorigenic mutations.

**Figure 2.**
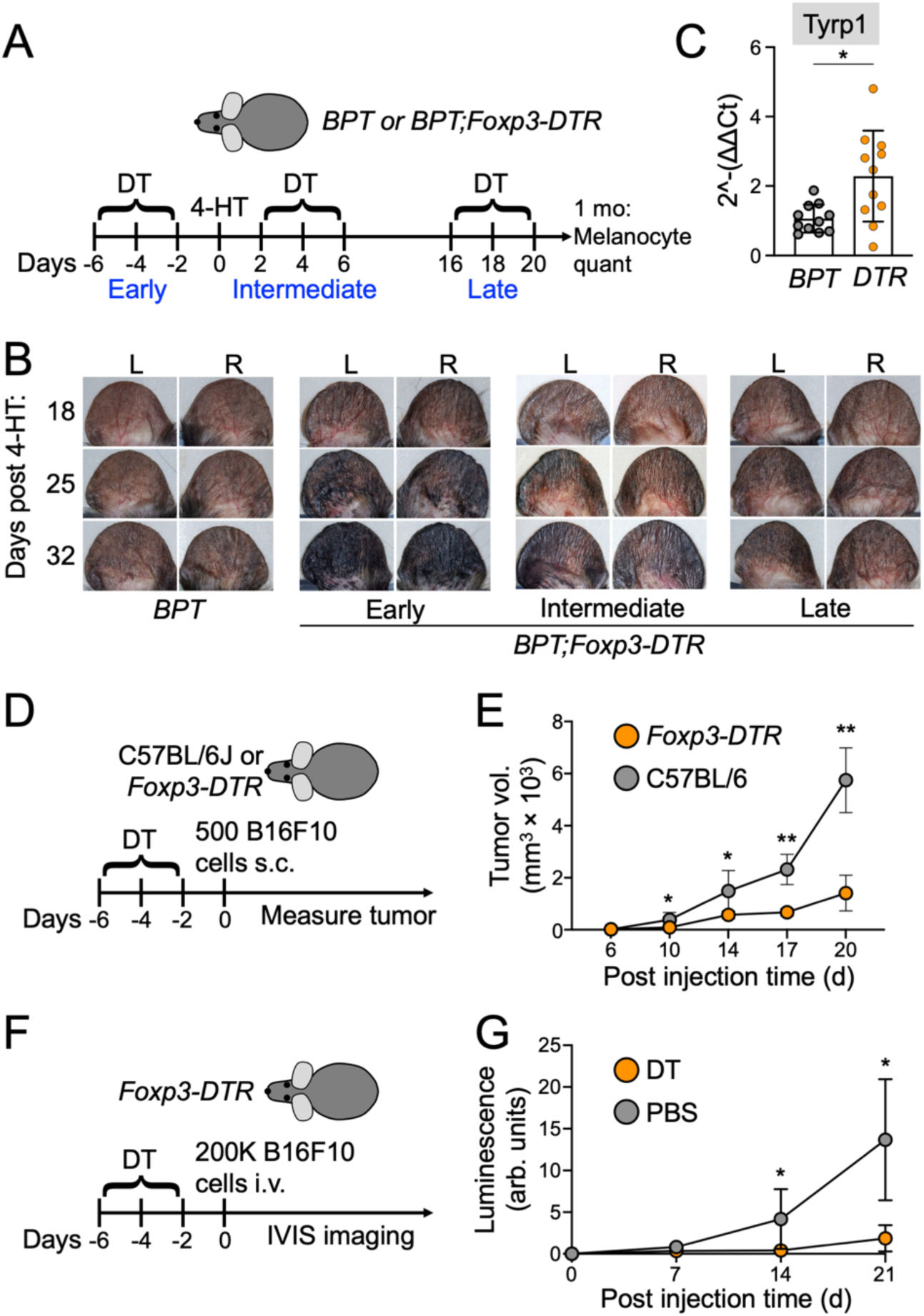
T_reg_ cell depletion promotes mutant melanocyte outgrowth in the *BPT* model. (A-B) *BPT* and *BPT;Foxp3-DTR* mice were subjected to DT treatment either 1 week before (early), just after (intermediate), or 2 weeks after (late) 4-HT painting, and melanocyte outgrowth assessed after 1 month. (A) Schematic of the experimental protocol. (B) Photographs showing representative melanocytic darkening at the indicated timepoints. (C) qRT-PCR quantification of *Tyrp1* expression in ear skin at the experimental endpoint. N = 11 mice per group. Error bars indicate SD. * denotes P ≤ 0.05, calculated by unpaired t-test. (D-E) *Foxp3-DTR* or control (C57BL/6J) mice were treated with DT and then injected s.c. with 500 B16F10 cells. Subsequent tumor growth was measured for three weeks. (D) Schematic diagram of the experimental approach. (E) Quantification of tumor growth, with error bars indicating SD. * and ** denote P ≤ 0.05 and P < 0.01, respectively, calculated by two-way ANOVA. N = 7 mice per group. (F-G) *Foxp3-DTR* mice were treated with DT or vehicle control (PBS), then injected i.v. with 2 × 10^5^ B16F10-Luciferase cells. Subsequent tumor growth was measured for three weeks by IVIS imaging. (F) Schematic diagram of the experimental approach. (G) Quantification of tumor growth, with error bars indicating SD. * denotes P ≤ 0.05, calculated by two-way ANOVA. N = 4 mice per group.

In light of these unexpected findings, we re-examined the role of T_reg_ cells in transplantable models of tumor growth. Subcutaneous (s.c.) implantation of syngeneic B16F10 melanoma cells is widely used to study anti-tumor immunity, and it has been applied previously to interrogate the effects of T_reg_ cells on tumor suppression^16,18,19^. To implement early stage T_reg_ cell depletion in the B16F10 system, *Foxp3-DTR* and wild type control mice were treated with the three dose DT regimen described above starting six days before B16F10 implantation (Fig. 2D). Transient T_reg_ cell deficiency significantly inhibited B16F10 tumor growth (Fig. 2E), confirming that T_reg_ cells constrain anti-tumor immunity in this model. In line with this interpretation, flow cytometric analysis of infiltrating immune cells in s.c. B16F10 tumors revealed a robust increase in both CD8^+^ and CD4^+^ T_conv_ cells in *Foxp3-DTR* animals (Fig. 2 - supplement 1C-D). Next, we performed an analogous set of experiments in mice receiving intravenous injections of B16F10, which leads to the formation of multifocal “metastatic” tumors in the lungs (Fig. 2F). T_reg_ cell depletion prior to B16F10 injection inhibited cancer outgrowth in this context (Fig. 2G), as well. We conclude that T_reg_ cells can both promote and antagonize incipient tumor formation in the skin, with the autochthonous *BPT* model revealing protective functions during early transformation that are either absent or substantially less important in transplantable B16F10 systems.

### T_reg_ cell depletion dramatically alters the cutaneous T cell-DC axis

To investigate how T_reg_ cell depletion promotes mutant melanocyte outgrowth in skin, we subjected *BPT;Foxp3-DTR;LSL-TdTomato* mice and *BPT;LSL-TdTomato* controls to early DT treatment (days -6, -4, and -2) and then assessed leukocyte content in the ear and draining lymph node (dLN) eight days after 4-HT painting (Fig. 3A, Fig. 3 - supplement 1A). T_reg_ cell depletion triggered a dramatic influx of multiple immune cell types into the skin, including T_conv_ cells, neutrophils (CD11b^+^Ly6G^+^), monocytes, macrophages, and DCs (CD11c^+^MHCII^+^F4/80^-^) (Fig. 3B-C). Skin T_conv_ cell expansion was apparent in both CD4^+^ and CD8^+^ compartments, but was much more obvious among CD4^+^ T_conv_ cells, which accounted for most of the cutaneous T cells at this time point (Fig. 3B). Among DCs, both cDC1 and cDC2 numbers increased, with cDC2s an order of magnitude more abundant than cDC1s in both T_reg_ cell deficient and T_reg_ cell sufficient skin (Fig. 3C). In the dLN, we observed expansion of all myeloid subsets, with more modest changes in T cell numbers (Fig. 3 - supplement 1B). T cell activation, however, as measured by upregulation of CD44 and downregulation of CD62L, was significantly enhanced in both CD4 and CD8 dLN subsets (Fig. 3 - supplement 1C). Upon restimulation with PMA/ionomycin *in vitro*, almost all dLN CD44^+^ CD4^+^ T_conv_ cells expressed IL-4, with little to no expression of IL-17 or IFNγ (Fig. 3 - supplement 1D). This observation is in line with prior work indicating that T_reg_ cells constrain an autoimmune T cell response with T_H_2 characteristics in the skin^48^.

**Figure 3.**
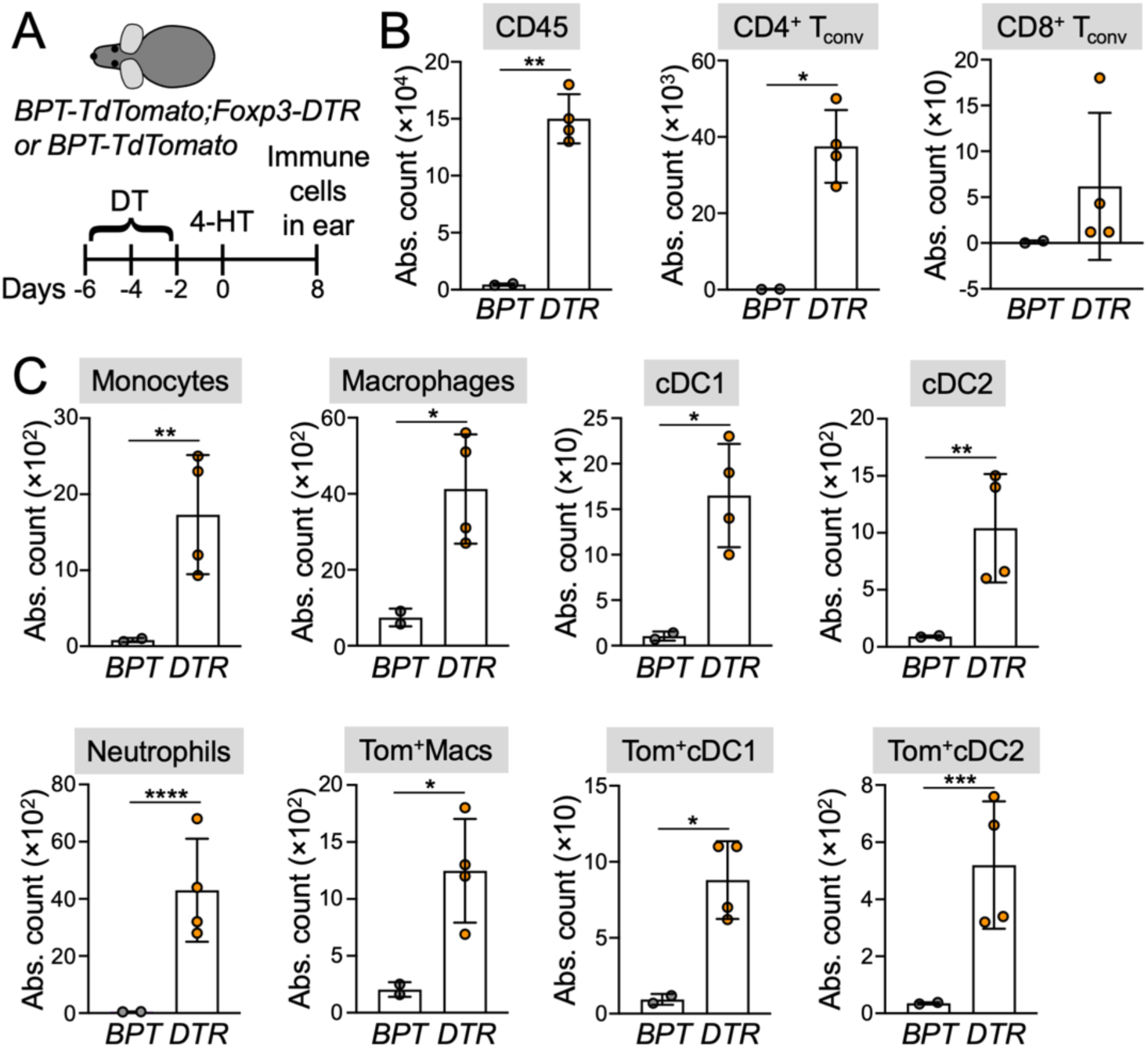
T_reg_ cell depletion drives multimodal inflammation of the skin. *BPT-TdTomato;Foxp3-DTR* mice and *BPT-TdTomato* controls were treated with DT, followed by 4-HT painting, and immune cells in the ear skin enumerated by flow cytometry after 8 days. (A) Schematic of the experimental protocol. (B) Quantification of the indicated T cell subsets. (C) Quantification of the indicated myeloid cell subsets. Tom^+^ = TdTomato positive. In B and C, *, **, ***, and **** denote P ≤ 0.05, P < 0.01, P < 0.001, and P < 0.0001, respectively, calculated by lognormal Welch’s t-test. N = 2 *BPT-TdTomato* mice and 4 *BPT-TdTomato;Foxp3-DTR* mice.

Next, we measured TdTomato fluorescence in each immune cell type to assess uptake of melanocyte antigen. Tumor sampling of this kind was largely restricted to macrophages, cDC1s, and cDC2s (Fig. 3C, Fig. 3 - supplement 1B). While T_reg_ cell depletion did not boost the fraction of TdTomato^+^ cells in each subset, their absolute numbers rose markedly due to increases in total macrophages and DCs. Notably, TdTomato^+^ DCs were observed in the dLN as well as the skin, indicative of successful antigen trafficking (Fig. 3 - supplement 1B). Taken together with the increased accumulation of cutaneous T_conv_ cells described above, these results are consistent with enhanced T_conv_ cell priming and activation in T_reg_ cell deficient skin, which is in line with prior reports^47–49^. What is more surprising is that this immune response failed to antagonize mutant melanocyte outgrowth in the *BPT* model.

### T_reg_ cell depletion induces the recruitment of tissue remodeling myeloid cells

To better understand how T_reg_ cells influence the skin microenvironment, we subjected *Foxp3-DTR* mice, along with C57BL/6J controls, to the three dose DT regimen and then profiled CD45^+^ cells in the ear by scRNA-seq eight days later (Fig. 4A). UMAP analysis of the resulting data revealed drastic differences in immune composition, encompassing both lymphoid and myeloid cells (Fig. 4B). T_reg_ cell depletion dramatically altered the cutaneous T cell compartment, as expected. Whereas control skin was largely devoid of T_conv_ cells, both CD4^+^ and CD8^+^ T_conv_ cells were readily apparent in T_reg_ cell-depleted skin (Fig. 4B, Fig. 4 - supplement 1A). *Gzmb* and *Fasl* were robustly expressed among CD8^+^ T_conv_ cells (Fig. 4 - supplement 1B), indicative of differentiation into armed cytotoxic T cells. CD4^+^ T_conv_ cells, for their part, exhibited high levels of *Gata3* and *Il4*, but low expression of *Rorc* and *Tbx21*, indicative of a T_H_2 response. *Cd69*, *Cd44*, and *Icos* were highly expressed across both subsets, consistent with ongoing T_conv_ cell activation. Collectively, these results suggest that T_reg_ cell depletion unleashes a strong T cell response in the skin, potentially targeting self-antigens or the cutaneous microbiota.

**Figure 4.**
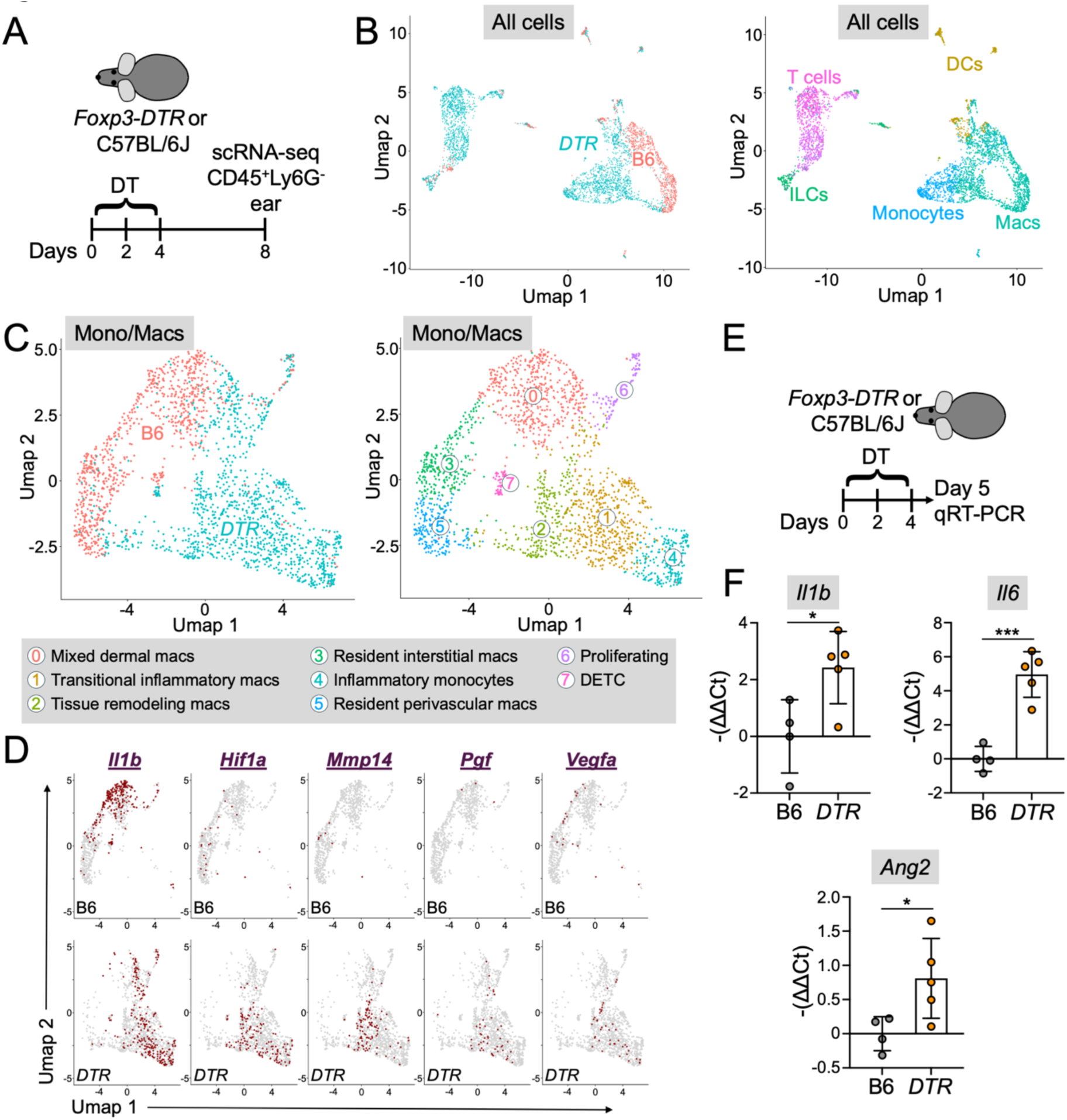
T_reg_ cell depletion leads to monocyte/macrophage inflammation and tissue remodeling. (A-D). CD45^+^ cells extracted from the ear skin of *Foxp3-DTR* mice or C57BL/6J controls were analyzed by scRNA-seq 8 days after DT treatment. (A) Schematic of the experimental protocol. (B) UMAP showing all sequenced cells from both samples, colored by sample on the left and by cell type on the right. (C) UMAP reclustering of the monocyte and macrophage clusters identified by the UMAP analysis of all cells in B. Cells are colored by sample on the left and by Seurat cluster on the right. The identities of individual Seurat clusters are shown in the legend below. (D) Feature plots of the monocyte/macrophage UMAP showing expression of the indicated inflammatory and tissue remodeling genes in the C57BL/6J (B6) and *Foxp3-DTR* (DTR) samples. (E-F) *Foxp3-DTR* mice and C57BL/6J controls were treated with DT and qRT-PCR performed on skin homogenates 5 days later. (E) Schematic of the experimental protocol. (F) Quantified expression of the indicated genes, with error bars denoting SD. *, **, and *** denote P ≤ 0.05, P < 0.01, and P < 0.001, respectively, calculated by unpaired t-test. N = 4 C57BL/6J mice and 5 *Foxp3-DTR* mice.

Seurat reclustering of Immgen-defined DCs revealed five clusters of cells whose fractional representations did not change substantially between C57BL/6J and *Foxp3-DTR* mice (Fig. 4 - supplement 1C). Although detailed analysis of individual genes revealed small increases in the expression of activation markers (*CD86* and *CD40*) and class II MHC (*H2-Ab1*) in certain DC clusters (Fig. 4 - supplement 1D), these transcriptional differences were quite modest overall. This was in sharp contrast to the marked increase in total cutaneous DC numbers we observed in T_reg_ cell deficient skin (Fig. 3). Taken together, these results indicate that T_reg_ cell depletion generates predominantly quantitative rather than to qualitative changes in the cutaneous DC compartment.

Conversely, the effects of T_reg_ cell deficiency among monocytes and macrophages were more striking, with several entirely new subsets appearing in the skin at the expense of other populations (Fig. 4C). UMAP replotting of the monocyte/macrophage clusters in the initial all-cell analysis identified two groups of resident cutaneous macrophages in C57BL/6J mice (Fig. 4C, Fig. 4 - supplement 2A). The first (DTR-M-cluster 3) expressed genes characteristic of resident macrophage identity (*Gpr34, Ms4a7, Pf4*), debris clearance (*Stab1, Cd93*), lipid processing (*Apoe*), and tissue maintenance (*Igf1*), implying a role in skin homeostasis. The second subset (DTR-M-cluster 5) exhibited indices of endothelial proximity (*Lyve1, Colec12*) and scavenging activity (*Cd209f, Cd209g, Colec12*), consistent with a resident perivascular macrophage identity.

T_reg_ cell depletion markedly diminished these subsets while inducing the emergence of three additional clusters. The first of these (DTR-M-cluster 4) bore transcriptional signatures characteristic of inflammatory monocytes (*Ly6a, Plac8, Sell, Trem3, S100a8*) and interferon signaling (*Ifitm6, Vcan, Isg15, Gbp2, Gbp4*). The second cluster (DTR-M-cluster 2) exhibited high levels of transcripts involved in macromolecular processing (*Lgmn, Fabp5*) and tissue remodeling (*Mmp14, Arg1*). This subset also expressed *Ccl24*, an established chemoattractant for eosinophils, which is consistent with the overall T_H_2 character of T_conv_ cells in T_reg_ cell-depleted skin. Importantly, DTR-M-cluster 2 cells also expressed weak but detectable levels of the interferon signaling module seen in DTR-M-cluster 4. Finally, the third cluster (DTR-M-cluster 1) exhibited strong interferon module expression and more modest levels of the inflammatory monocyte and tissue remodeling modules. Taken together, these results were consistent with the ongoing recruitment of inflammatory monocytes (DTR-M-cluster 4) into T_reg_ cell depleted skin and their differentiation into tissue remodeling macrophages (DTR-M-cluster 2) via an inflammatory transitional macrophage subset (DTR-M-cluster 1). Indeed, pseudotime analysis identified a predominant differentiation trajectory starting at DTR-M-cluster 4 and moving sequentially through cluster 1 and then cluster 2 (Fig. 4 - supplement 2B-C). Interestingly, this trajectory then connected cluster 2 with DTR-M-cluster 0, a macrophage population expressing indices of tissue remodeling (*Mmp12*), lipid metabolism and alternative activation (*Ramp3, Slc27a3*), and immunoregulation (*Cd226, Klrb1b*). Cluster 0 also expressed skin epithelial markers (*Skint3, Hepacam2*), consistent with skin residence or the recent uptake of material from skin resident cells. Notably, cluster 0 was well represented in both *Foxp3-DTR* and C57BL/6J skin, implying that it could be a more long-lived, terminal differentiation state for monocyte-derived macrophages.

Further analysis of the scRNA-seq data revealed elevated expression of stromal remodeling genes, including the hypoxia-induced transcription factor *Hif1a* and the matrix metalloprotease *Mmp14*, by the three monocyte derived subsets (DTR-M-clusters 1, 2, and 4) that emerge in the context of T_reg_ cell deficiency (Fig. 4D). These subsets also displayed increased levels of the angiogenic factors *Vegfa* and *Pgf*, which were only weakly expressed by resident macrophages (DTR-M-clusters 3 and 5). The inflammatory cytokine *Il1b* was robustly expressed by the inflammatory subsets and also by the mixed identity macrophage cluster (DTR-M-cluster 0). This expression pattern would be expected to translate into higher overall *Il1b* production after T_reg_ cell depletion, given the dramatic quantitative increase in cutaneous monocytes and macrophages we observed under these conditions (Fig. 3).

To confirm and extend these results, we subjected *Foxp3-DTR* mice and C57BL/6J controls to the three dose DT regimen and then measured the expression of key inflammatory factors by qRT-PCR five days later (Fig. 4E). T_reg_ cell depletion triggered strong upregulation of both *Il1b* and a second inflammatory cytokine, *Il6* (Fig. 4F). We also documented increased expression of *Ang2*, which encodes a secreted factor that destabilizes the vasculature (Fig. 4F). Taken together with the scRNA-seq data described above, these results document a broad and rapid myeloid inflammatory response in T_reg_ cell deficient skin.

### T_reg_ cells maintain vascular homeostasis during premalignant melanomagenesis

The upregulation of angiogenic and blood vessel remodeling factors that we observed in T_reg_ cell deficient skin suggested that these cells might promote vascular stability. To evaluate this hypothesis, we used two photon microscopy to compare cutaneous blood vessel architecture in T_reg_ cell depleted mice with that of healthy controls five days after DT treatment (Fig. 5A). In normal C57BL/6 skin, i.v. injected high molecular weight fluorescent dextran was largely constrained within the vasculature. In *Foxp3-DTR* mice, however, a substantial amount of injected dextran leaked into the surrounding dermal interstitium (Fig. 5B), indicative of vascular permeability. To quantify this vessel destabilization phenotype, we calculated the fraction of dextran-positive area using Z-projection images of the skin. This approach revealed a nearly two-fold increase in the scope of dextran diffusion in T_reg_ cell deficient skin, consistent with a role for these cells in maintaining vascular integrity (Fig. 5C). As an alternative quantification approach, we i.v. injected the mice with the vital dye Evans Blue and then assessed its dissemination into extravascular ear tissue 24 hours later (Fig. 5D). We again found that T_reg_ cell depletion induced a significant, 2-3 fold increase in blood vessel permeability (Fig. 5E).

**Figure 5.**
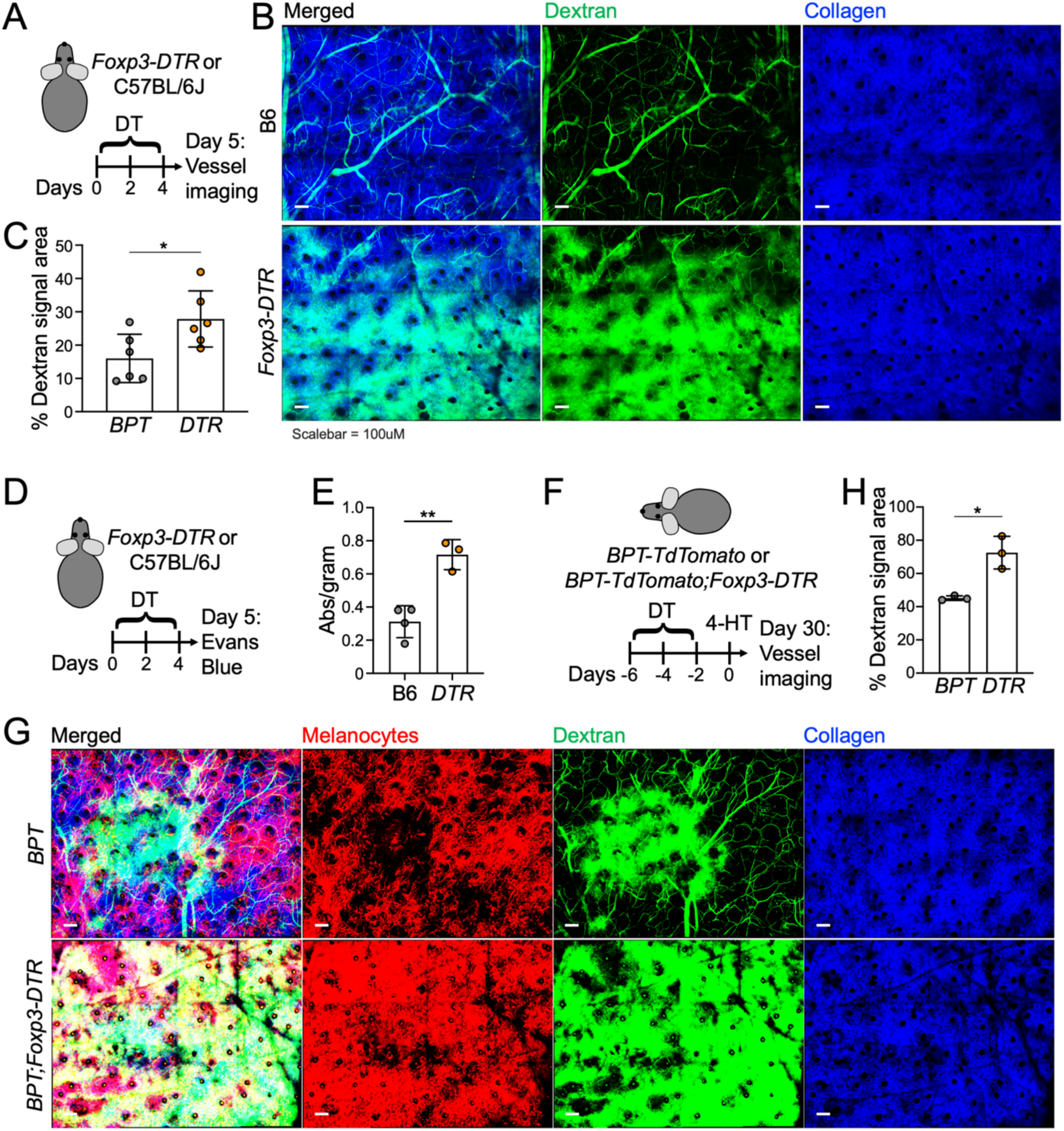
T_reg_ cell depletion destabilizes vasculature in the skin. (A-C) *Foxp3-DTR* mice and C57BL/6J controls were treated with DT and cutaneous vasculature imaged 5 days later by two-photon microscopy after dextran injection. (A) Schematic of the experimental protocol. (B) Representative images of dextran and dermal collagen in C57BL/6J (top) and *Foxp3-DTR* (bottom) skin. Scale bars = 100 μm. (C) Quantification of vascular leakage by fractional area occupied by fluorescent dextran. N = 6 mice per group. (D-E) *Foxp3-DTR* mice and C57BL/6J controls were treated with DT and vasculature leakage in the ears assessed 5 days later by Evans Blue injection. (D) Schematic of the experimental protocol. (E) Quantification of Evans Blue leakage into the skin. N = 4 C57BL/6J (B6) mice and 3 *Foxp3-DTR* mice. (F-H) *BPT-TdTomato* and *BPT-TdTomato;Foxp3-DTR* mice were treated with DT, painted with 4-HT, and then subjected to two-photon imaging after dextran injection 30 days later. (F) Schematic of the experimental protocol. (G) Representative images of dextran, dermal collagen, and TdTomato^+^ melanocytes in *BPT-TdTomato* (top) and *BPT-TdTomato;Foxp3-DTR* (bottom) skin. Scale bars = 100 μm. (H) Quantification of vascular leakage by fractional area occupied by fluorescent dextran. N = 3 mice per group. All error bars indicate SD. * and ** denote P ≤ 0.05 and P < 0.01, respectively, calculated by unpaired t-test.

Next, we examined the interplay between vascular integrity and T_reg_ cells in the context of early tumorigenesis. *BPT*;*LSL-TdTomato;Foxp3-DTR* mice and *BPT*;*LSL-TdTomato* controls were subjected to early DT treatment, 4-HT painting, and two-photon imaging with blood vessel tracing performed 30 days after oncogene induction (Figure 5G). TdTomato^+^ melanocyte expansion was readily apparent in both experimental groups, although more pronounced in *BPT*;*LSL-TdTomato;Foxp3-DTR* animals, as expected. Importantly, the increased melanocyte load in T_reg_ cell deficient skin was accompanied by severe dextran leakage, which obscured vessel architecture in large regions of tissue (Fig. 5G-H). These results strongly suggest that T_reg_ cells and the inflammatory responses they keep in check control vascular integrity in the premalignant microenvironment.

### UVB radiation drives cutaneous inflammation and premalignant melanocyte expansion

The link between cutaneous inflammation and melanoproliferation that we observed in the context of T_reg_ cell deficiency prompted us to investigate whether other inflammatory stimuli might affect the premalignant microenvironment in similar ways. UV light is the major environmental trigger of human melanoma^38^. Beyond causing DNA damage, UV exposure also induces a potent inflammatory response characterized by macrophage activation, the upregulation of pro-inflammatory mediators, such as IL-1β and TNF ^24,50–54^, and VEGF production, which contributes to a pro-angiogenic environment^55,56^. Given the similarities between these processes and the effects of T_reg_ cell depletion in the skin, we hypothesized that UV exposure would accelerate tumor progression via analogous mechanisms. To this end, we irradiated the ear skin of *BPT* mice with 2 kJ/m² UVB three days after 4-HT treatment (Fig. 6A). This dose is known to accelerate melanoma progression in autochthonous models, and while it does induce DNA mutations, driver mutations causative for the outgrowth phenotype have not been identified^37^. UVB markedly enhanced the expansion of mutant melanocytes, leading to increased ear darkening and elevated *Tyrp1* expression within a month of oncogene induction (Fig. 6B-C). This outgrowth essentially mirrored the consequences of T_reg_ cell deficiency.

**Figure 6.**
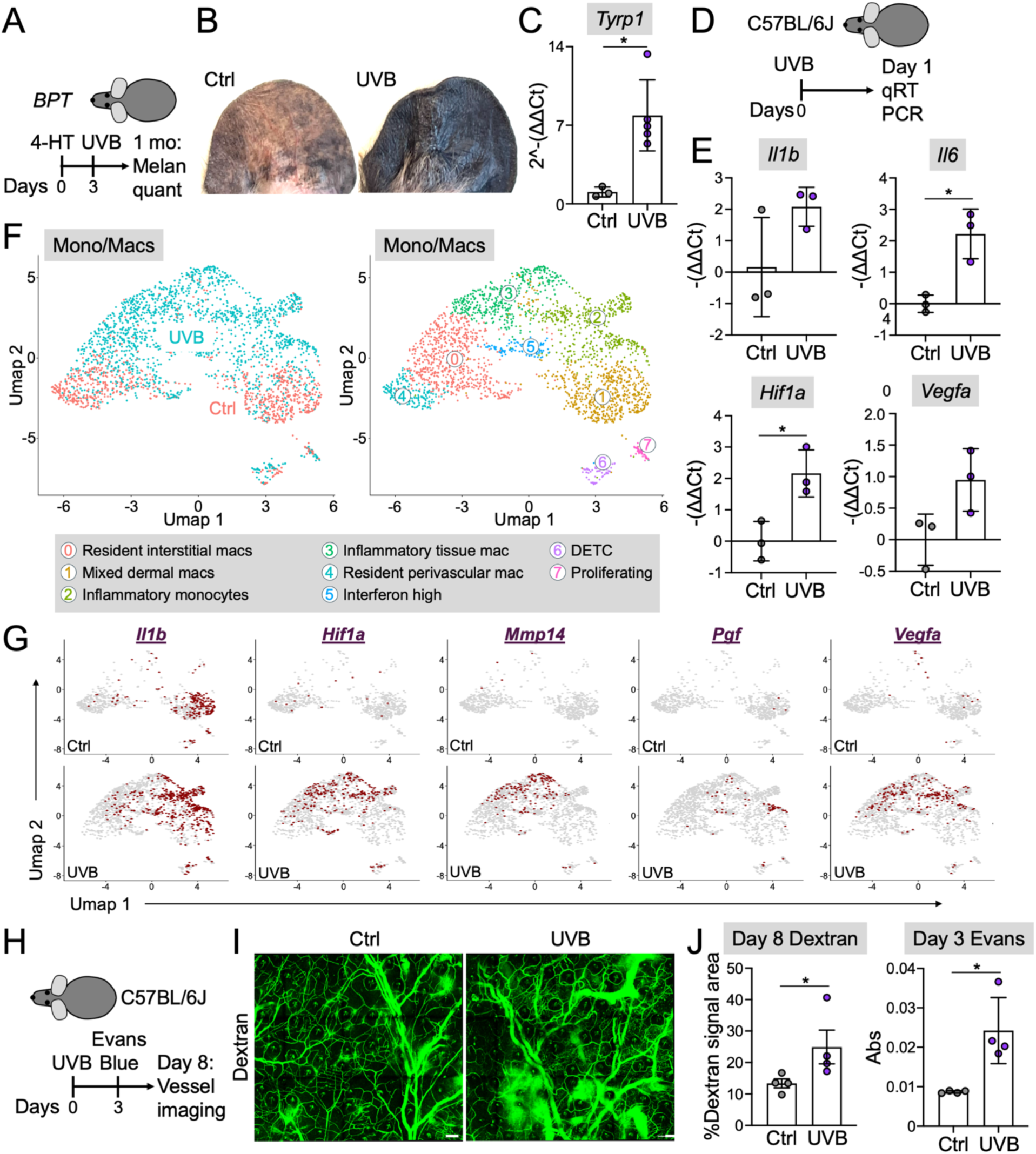
UVB irradiation drives myeloid inflammation and tissue remodeling in the skin. (A-B) *BPT* mice were UVB irradiated 3 days after 4-HT painting and melanocyte outgrowth assessed at the 30 day timepoint. (A) Schematic of the experimental protocol. (B) Photographs showing representative melanocytic darkening in unirradiated (Ctrl) and UVB-irradiated mice. (C) qRT-PCR quantification of *Tyrp1* expression in ear skin at day 30. N = 3 Ctrl mice and 5 UVB-treated mice. Error bars indicate SD. * denotes P ≤ 0.05, calculated by unpaired t-test. (D-E) C57BL/6J mice were UVB- or mock-irradiated and qRT-PCR performed on skin homogenates one day later. (D) Schematic of the experimental protocol. (E) Quantified expression of the indicated genes. N = 3 mice per group. (F-G) C57BL/6J mice were exposed to UVB or mock irradiation and CD45^+^ cells from the ear skin analyzed by scRNA-seq 7 days later. (F) UMAP reclustering of the monocyte and macrophage clusters identified by UMAP analysis of all cells (see Fig. 6 - supplement 2). Cells are colored by sample on the left, and by Seurat cluster on the right. The identities of individual Seurat clusters are shown in the legend below. (G) Feature plots of the monocyte/macrophage UMAP showing expression of the indicated inflammatory and tissue remodeling genes in the Ctrl and UVB samples. (H-J) C57BL/6J mice were UVB- or mock-irradiated and cutaneous vasculature integrity assessed 3 days later by Evans Blue injection or 8 days later by two-photon microscopy after dextran injection. (H) Schematic of the experimental protocol. (I) Representative images of dextran in mock- (left) and UVB-irradiated (right) skin. Scale bars = 100 μm. (J) Left, vascular leakage at 8 days post UVB, quantified by fractional area occupied by fluorescent dextran. N = 4 mice per group. Right, quantification of Evans Blue leakage into the skin. N = 4 mice per group. All error bars indicate SD. * denotes P ≤ 0.05, calculated by unpaired t-test.

To better understand the microenvironmental effects of UVB in this context, we used qRT-PCR to quantify the expression of inflammatory and angiogenic factors in wild type skin after UV irradiation (Fig. 6D, Fig. 6 - supplement 1A). Within 24 hours of UVB treatment, we observed upregulation of both *Il1b* and *Il6*, along with increased expression of *Vegfa* and *Hif1a* (Fig. 6E). Expression levels of *Il1b* and *Il6* remained high seven days post-UVB (Fig. 6 - supplement 1B). These results indicated that UVB induces a rapid inflammatory and tissue remodeling response in the skin with molecular properties that are similar to the effects of T_reg_ cell depletion. To investigate the cellular basis for this response in more detail, we profiled immune infiltrates in the skin by flow cytometry seven days after UVB treatment (Fig. 6 - supplement 1C). Neutrophil, monocyte, macrophage, and DC levels were all markedly enhanced by UVB (Fig. 6 - supplement 1D). CD4^+^ T_conv_ cell and T_reg_ cell numbers also increased, which was consistent with prior reports^42,43,57,58^, while CD8^+^ T_conv_ cells remained unchanged (Fig. 6 - supplement 1E). Hence, whereas certain aspects of UVB-induced inflammation mirrored those of T_reg_ cell-depleted skin, other features, most notably the presence of T_reg_ cells themselves, were distinct.

Next, we performed scRNA-seq to profile the diversity of immune cells (CD45^+^) in the ear skin seven days after 2 kJ/m² UVB irradiation. These cells were predominantly of myeloid lineage, consistent with the flow cytometric results described above (Fig. 6 - supplement 2A-B). The most striking UV-induced changes manifested in the monocyte/macrophage compartment, where two entirely new subsets appeared in irradiated skin (Fig. 6F, Fig. 6 - supplement 2C). These subsets bore a striking resemblance to the monocyte derived populations observed in T_reg_ cell deficient skin (Fig. 4). The first (UV-M-cluster 2) strongly expressed transcripts indicative of inflammatory monocyte identity (*Trem1, Plac8*) and interferon signaling (*Vcan, Ifitm1, Ifitm6*) and more weakly expressed transcripts associated with tissue remodeling (*Spp1, Gpnmb, Mmp14*) and lipid handling (*Fabp5, Apoc2, Hilpda*). This pattern was reversed in the second subset (UV-M-cluster 3), which expressed high levels of the remodeling and lipid handling modules and lower levels of the monocyte and interferon signatures.

Taken together, these results were consistent with UVB-induced recruitment of inflammatory monocytes from the blood, followed by their differentiation into inflammatory, tissue remodeling macrophages. Importantly, *Il1b*, *Mmp14*, *Hif1a*, *Vegfa*, and *Pgf* expression was strong across all UVB-induced subsets (Fig. 6G), suggestive of increased inflammatory and angiogenic activity. To assess this hypothesis directly, we performed intravital blood vessel tracing of ear skin and Evans Blue assays, eight and three days after UVB exposure, respectively (Fig. 6H). These experiments revealed significantly more blood vessel leakage in irradiated ear skin relative to mock-irradiated controls (Fig. 6I-J), consistent with the interpretation that UVB disrupts the cutaneous vasculature. Hence, while UVB-induced skin inflammation differs somewhat from the response to T_reg_ cell depletion, both feature the infiltration of tissue remodeling macrophages and vascular instability.

### Contact dermatitis is sufficient to drive dysregulated melanocyte outgrowth

Having demonstrated that an environmental irritant with known immunomodulatory properties (UVB) could induce dysregulated melanocyte outgrowth, we next asked whether a topically applied chemical irritant would have similar effects. 2,4-dinitrofluorobenzene (DNFB) drives potent cutaneous inflammatory responses via covalent modification of skin proteins and other biomolecules, leading to oxidative stress, adaptive immune priming, and type 4 hypersensitivity^59^. We were particularly interested in DNFB because of prior work indicating that it could strongly induce IL-1β and IL-6, a feature we had observed in both UVB-treated and T_reg_ cell deficient skin.

To induce DNFB dependent inflammation, we employed a classical approach in which mice are first sensitized with DNFB on their abdomen and then challenged on one of their ears five days later (Fig. 7A, Fig. 7 - supplement 1A). This protocol induced substantial swelling on the DNFB-treated but not the contralateral, vehicle-treated ear (Fig. 7 - supplement 1B), indicative of strong, DNFB-dependent type 4 hypersensitivity. Using qRT-PCR, we observed dramatic upregulation of *Il1b*, *Il6*, *Hif1a*, and *Ang2* following DNFB elicitation (Fig. 7B). DNFB elicitation also induced marked vascular leakage, as measured by Evans Blue staining (Fig. 7C). Using flow cytometry, we documented a dramatic increase in immune cellularity within two days of DNFB elicitation, encompassing neutrophils, DCs, monocytes, macrophages, and both CD4^+^ and CD8^+^ T_conv_ cells (Fig. 7 - supplement 1C-D). Hence, DNFB-induced inflammation recapitulated key characteristics of the immunological responses seen in UVB-treated and T_reg_ cell-depleted skin.

**Figure 7.**
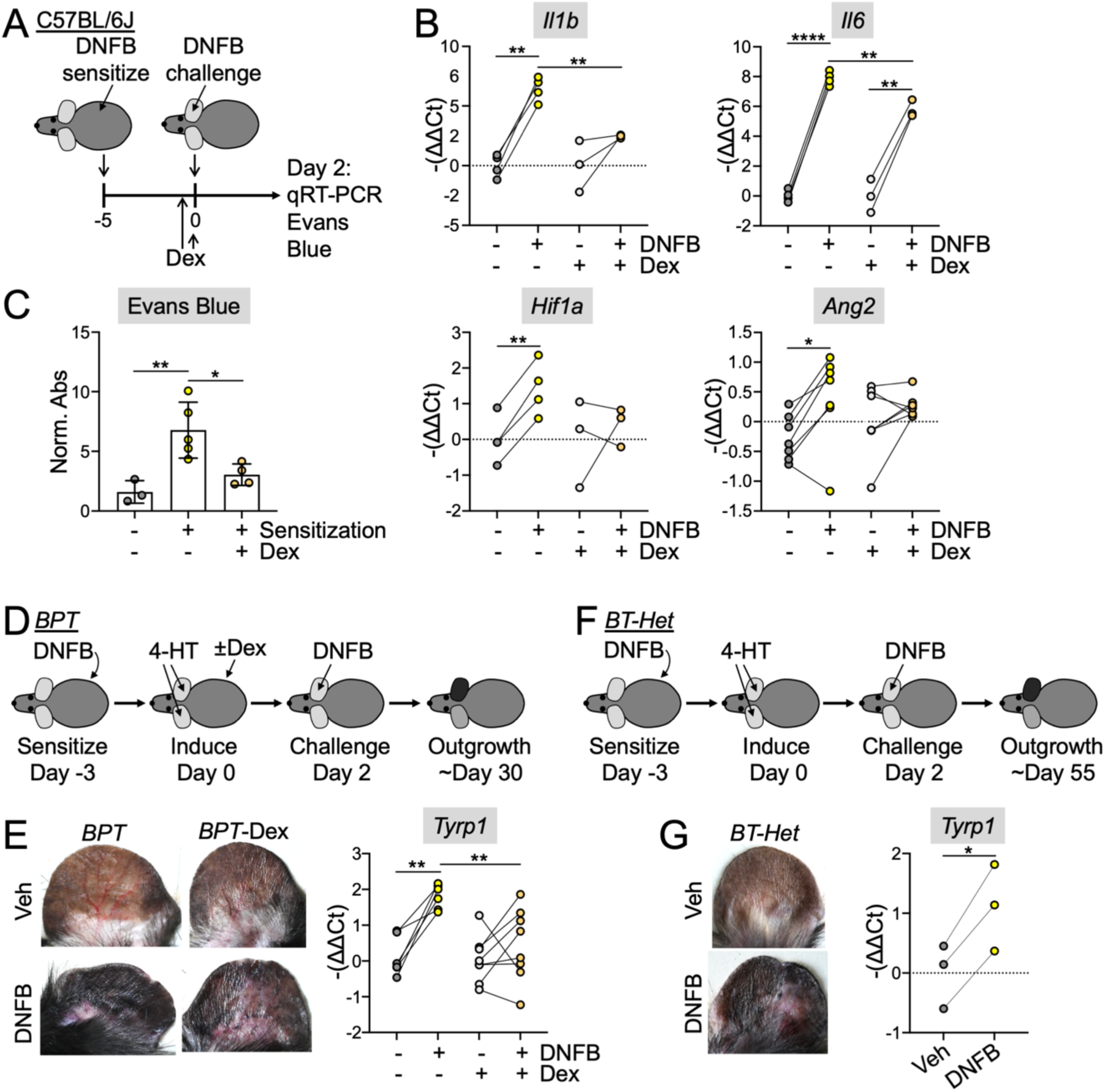
Contact hypersensitivity drives inflammation dependent melanocyte expansion. (A-C) C57BL/6J mice were sensitized with DNFB and then rechallenged on one ear in the presence or absence of Dex therapy. qRT-PCR analysis of inflammatory genes and Evans Blue analysis of vascular leakage was performed 2 days after elicitation. (A) Schematic of the experimental protocol. (B) Expression of the indicated inflammatory genes was quantified for the indicated DNFB and Dex treatment conditions. N ≥ 3 mice per group. ** and **** denote P < 0.01 and P < 0.0001, respectively, calculated by paired t-test for +/- DNFB comparisons and unpaired t-test for +/- Dex comparisons. (C) Quantification of Evans Blue leakage into the skin. N ≥ 3 mice per group. * and ** denote P ≤ 0.05 and P < 0.01, respectively, calculated by one-way ANOVA. Error bars indicate SD. (D-E) *BPT* mice were sensitized to DNFB, 4-HT-painted on both ears, and then rechallenged with DNFB on one ear in the presence or absence of Dex therapy. Melanocyte outgrowth was quantified photographically and by qRT-PCR 30 days later. (D) Schematic of the experimental protocol. (E) Left, photographs showing representative melanocytic darkening in DNFB- and vehicle-treated ears, with elicitation performed in the presence or absence of Dex. Right, qRT-PCR quantification of *Tyrp1* expression in ear skin at the experimental endpoint. N ≥ 6 mice per group. ** denotes P < 0.01, calculated by paired t-test for +/- DNFB comparisons and unpaired t-test for +/- Dex comparisons. (F-G) *BPT-Het* mice were sensitized to DNFB, 4-HT-painted on both ears, and then rechallenged with DNFB on one ear. Melanocyte outgrowth was quantified photographically and by qRT-PCR 55 days later. (F) Schematic of the experimental protocol. (G) Left, photographs showing representative melanocytic darkening in DNFB- and vehicle-treated ears. Right, qRT-PCR quantification of *Tyrp1* expression in ear skin at the experimental endpoint. N = 3 mice per group. * denotes P ≤ 0.05, calculated by paired t-test.

Next, we assessed the capacity of DNFB-induced type 4 inflammation to promote mutant melanocyte outgrowth. BPT mice were sensitized with DNFB, painted with 4-HT on both ears three days later, and then challenged with DNFB on one ear two days after that (Fig. 7D). DNFB-treated ears exhibited substantially more darkening and *Tyrp1* expression than their contralateral counterparts (Fig. 7E), indicative of profoundly dysregulated melanocyte expansion. This DNFB induced effect was similar to but more robust than what we had observed in T_reg_ cell-depleted or UVB-treated mice, prompting us to speculate that it might be capable of promoting melanocyte outgrowth in non-oncogenic conditions. To test this hypothesis, we applied the same experimental protocol to *LSL-Braf^V600E^*;*Pten^fl/+^;Tyr::CreERT2* (*BT-Het)* mice, which retain one copy of *Pten* (Fig. 7F). *BT-Het* mice fail to develop melanoma after 4-HT treatment, with melanocyte expansion arresting at the nevus stage^45^. DNFB elicitation dramatically accelerated melanocyte outgrowth in these animals (Fig. 7G), suggesting that cutaneous inflammation can drive melanoproliferation even in the absence of tumorigenesis.

Finally, to determine whether cutaneous inflammation was the actual cause of DNFB elicitation-induced melanoproliferation, we treated 4-HT- and DNFB-painted *BPT* mice with dexamethasone (Dex), a powerful immunosuppressive anti-inflammatory drug. Dex was administered both by i.p. injection one hour before DNFB elicitation and in a topical manner concomitantly with DNFB (Fig. 7A, 7D, S7A). In the context of Dex treatment, we observed marked reductions in *Il1b*, *Hif1a*, and *Ang2* expression in DNFB-painted ears (Fig. 7B), which were also less swollen than the corresponding ears on mice receiving no Dex (Fig. 7 - supplement 1B). Interestingly, *Il6* levels were only modestly inhibited by Dex (Fig. 7B). Notwithstanding this mixed effect on inflammatory indices, we observed a substantial reduction in DNFB elicitation-induced melanocyte outgrowth in Dex-treated mice (Fig. 7E), which was accompanied by marked attenuation of the vascular leakage phenotype (Fig. 7C). Taken together, these data demonstrate that type 4 inflammation can drive robust outgrowth of melanocytes, and they also suggest that blood vessel remodeling plays a particularly important role in this process.

## Discussion

The effects of acute inflammation on the premalignant phase of cancer progression are poorly understood, due in no small part to difficulties in studying early disease using standard transplantable tumor models. To address this issue, we combined the autochthonous *BPT* model with three inflammatory triggers: T_reg_ cell depletion, UVB irradiation, and type 4 hypersensitivity. Strikingly, all three perturbations markedly enhanced the outgrowth of mutant melanocytes, despite the fact that they are known to elicit cutaneous inflammation in mechanistically distinct ways. T_reg_ cell depletion drives systemic T_conv_ cell activation, leading to extensive infiltration of CD8^+^ T_conv_ cells, CD4^+^ T_H_2 effector cells, and inflammatory myeloid cells into the skin^47,48,60^. DNFB-induced hypersensitivity also drives T_conv_ cell and myeloid inflammation, but the response is more T_H_1-skewed and constrained to the skin itself^61^. In contrast, UVB promotes robust cutaneous recruitment of myeloid cells along with CD4^+^ T_conv_ and T_reg_ cells, but not CD8^+^ T_conv_ cells^24,25,42,44^. Although the distinct features of these three responses were readily apparent in our experiments, certain critical attributes were shared, including the robust accumulation of inflammatory monocytes and macrophages, increased expression of *Il1b* and *Il6*, and the destabilization of cutaneous vasculature, accompanied by the upregulation of tissue remodeling and angiogenesis programs. *Hif1a*, *Vegfa*, and *Ang2* likely function together within this axis to drive vascular remodeling, promoting vessel destabilization, angiogenesis, and increased permeability. Importantly, reversal of inflammatory tissue remodeling with Dex inhibited DNFB-induced outgrowth of mutant melanocytes. Vascular destabilization and angiogenesis are thought to promote the progression of small tumors into more aggressive and metastatic states^62^. Our results indicate that this “angiogenic switch” may be initiated and/or activated at a much earlier, premalignant stage.

T_reg_ cells are widely thought to facilitate tumor growth by restraining T_conv_ cell responses against tumor-specific antigens^63^. Our findings challenge this view by showing that T_reg_ cell depletion can enhance the outgrowth of mutant melanocytes during the early, premalignant phase of disease progression. This unexpected phenotype is likely caused by dysregulated T_conv_ cell responses against non-melanocytic self-antigens and/or cutaneous microbiota, leading to tissue damage, inflammatory myeloid infiltration, and the tissue remodeling response described above. Additional studies will be required to establish the precise chain of causation. At this stage, however, it seems fair to conclude that T_reg_ cells can exert both pro- and anti-tumorigenic effects on melanoma, depending on the stage of disease progression and on specific microenvironmental features that emerge through the course of tumor growth. In light of our findings, immunotherapeutic efforts centered on the modulation of T_reg_ cell activity should be evaluated with this more holistic conception of T_reg_ cell function in mind.

The inhibition of melanocyte outgrowth by T_reg_ cells in the *BPT* model is particularly striking given prior results from other labs (and our own data) showing that T_reg_ cells play a protumorigenic role in transplantable models of disease^16–19^. Although transplantable systems are easier to implement and quantify, the injection of a bolus of cultured cancer cells, often with additional components like matrigel, disrupts the tissue interactions and stromal-immune crosstalk that define the microenvironment of nascent tumors. The capacity of autochthonous models to recapitulate these microenvironmental features is therefore critical for accurately representing the early, premalignant stage of disease.

It is becoming increasingly clear that UV light can promote melanocyte proliferation and melanoma independently of its capacity to induce oncogenic mutations. We and others have shown that a single dose of UVB is sufficient to promote melanomagenesis and that timing (UVB delivered immediately before or during the formation of driver oncogenes but not after) is critical for generating melanomagenic effects^31,37,39–41^. Taken together, these findings suggest that UV enhances melanoma outgrowth in part by altering the local tumor microenvironment. Our present results lend further support to this idea by closely tying UVB-induced melanocyte expansion to an inflammatory immune response in the skin, which is consistent with prior work showing that acute UVB irradiation promotes the expansion of both oncogenically mutated and unmutated melanocytes in an inflammation dependent manner^64–66^. Interestingly, two of these studies directly implicated inflammatory macrophages in the melanocyte proliferation response^65,66^. Identifying the UVB-elicited factors underlying myeloid recruitment and also the molecular pathways by which these cells activate melanocytes could reveal new translational avenues for mitigating the oncogenic effects of UV. Notably, UVB-induced inflammation also leads to local T_reg_ cell expansion in the skin, which could facilitate premalignant outgrowth by restraining melanocyte-reactive T_conv_ cell activity. Although the relative importance of myeloid cells *vis-a-vis* T_reg_ cells in this context remains to be determined, our observations that T_reg_ cell depletion and DNFB-induced hypersensitivity both enhance early melanocyte expansion while simultaneously promoting T_conv_ cell activation and infiltration implies a more critical role for myeloid cells in this context.

Regular aspirin use has been associated with reduced melanoma incidence in postmenopausal women^67^, implying that anti-inflammatory drugs could potentially be used as a chemoprevention strategy for this disease. While clinical findings like this must be interpreted cautiously, they are consistent with the causal role for inflammatory signaling in early melanomagenesis that our results support. That Dex suppresses DNFB-induced melanocyte outgrowth in our model provides a direct experimental correlate and motivates more targeted investigation of specific anti-inflammatory mediators as potential preventive agents in individuals with elevated melanoma risk.

Our findings that contact hypersensitivity can promote the expansion of non-malignant *BT-Het* melanocytes have interesting implications for benign hyperpigmentation conditions, such as post-inflammatory hyperpigmentation, Riehl’s melanosis, and melasma^68–74^. These disorders can lead to significant psychosocial stress, and billions of dollars are spent annually on treatments to reduce excess pigmentation. Post-inflammatory hyperpigmentation and Riehl’s melanosis usually occur following T_conv_ cell dependent inflammatory conditions such as atopic dermatitis or allergic contact dermatitis, whereas melasma is associated with exuberant mast cell and T_H_2 inflammation. In each case, the condition is defined by increased proliferation of melanocytes leading to increased epidermal pigmentation. In light of our results, it is tempting to speculate that a better understanding of *BT-Het* melanocyte outgrowth in the context of cutaneous inflammation could inform the treatment of both disorders.

While chronic inflammation is an established hallmark of cancer, acute inflammation is often presented as a critical component of anti-tumor immunity. Our results indicate that the cancer cell extrinsic consequences of acute inflammation can have the opposite effect by laying a stromal foundation for premalignant expansion. Whether pro- or anti-tumoral effects predominate in a given situation will likely depend on contextual features such as the structure of the tissue in question, the size and stage of the tumor, and the tone of local immune cells. Identifying these inflammatory determinants could facilitate improved cancer prevention and immunotherapy.

## Methods

### Mice

The animal protocols used in this study were approved by the Institutional Animal Care and Use Committee of Memorial Sloan Kettering Cancer Center. Unless specified, experimental mice were 8-10 weeks old. All comparative analyses were performed on age- and sex-matched cohorts, which were either male or female. *Foxp3-DTR* mice were kindly provided by A. Y. Rudensky. *BPT* (B6.Cg-Tg(Tyr-cre/ERT2)13Bos;Braf-tm1Mmcm;Pten-tm1Hwu/BosJ), C57BL/6J, *Cd11c-yfp* (B6.Cg-Tg(Itgax-Venus)1Mnz/J), and *LSL-TdTomato* (B6.Cg-Gt(ROSA)26Sor-tm9(CAG-tdTomato)Hze/J) mice were purchased from the Jackson Laboratory. All reported strains and combinations thereof were bred and maintained at Memorial Sloan-Kettering Cancer Center (MSKCC).

### T_reg_ Cell Depletion

To achieve sustained T_reg_ cell depletion, a total of three intraperitoneal (i.p.) DT (25 mg/kg) injections were administered to *Foxp3-DTR* mice at 48-hour intervals. T_reg_ cell clearance was verified by flow cytometric quantification of Foxp3^+^ T cells in the blood.

### UVB irradiation

Mice were anesthetized with isoflurane and laid prone on a warming blanket. They were then irradiated with UVB using a Daavlin hand-held broad band (280-320nm filter with sharp cutoff) lamp (5.2 mW/s). The lamp was metered before each irradiation to ensure consistency and distance from mice was maintained at 0 cm. Mock irradiation was conducted using a standard incandescent desk lamp which was confirmed to not emit UVB.

### Subcutaneous B16F10 Implantation

500 B16F10 cells were resuspended in RPMI and mixed in a 3:1 ratio with growth factor–reduced matrigel. The sample was then subcutaneously (s.c.) injected in the flank of recipient mice. Subsequent primary tumor outgrowth was monitored by measuring tumor length (L) and width (W). Volume was then calculated as πLW2/6.

### Cell Implantation for Metastasis

200,000 of luciferase expressing B16F10 cells were injected intravenously (i.v., tail vein) into recipient mice. Tumor growth was monitored weekly thereafter by bioluminescent (IVIS) imaging.

Prior to imaging, mice were anesthetized with isoflurane and their thoracoabdominal fur shaved using an electric clipper. D-luciferin (150 mg/kg) was injected retro-orbitally into each mouse, and bioluminescence recorded using an IVIS Spectrum Xenogen instrument (Caliper Life Sciences) equipped with Living Image Software v.2.50. A head-to-tail-base ROI (region of interest) of uniform size was applied to all subjects to measure the total flux (photons/second) of metastatic cancer cells.

### 4-HT Application

Mice were placed under isoflurane anesthesia, and 32 μg of 4-Hydroxytamoxifen (Sigma, H6278) in DMSO was delivered via micropipette to each ear. Isoflurane was maintained for 15-20 minutes to ensure unhindered absorption of the drug.

### DNFB Sensitization and Elicitation

Mice were placed under isoflurane anesthesia, and 25 μL of 0.5% DNFB (1-fluoro-2,4-dinitrobenzene) in 1:4 olive oil:acetone was applied via micropipette to the shaved abdomen. The treated area was allowed to dry for 4 minutes. Hindleg nails were clipped to prevent subsequent scratching of the sensitized site (abdomen). 5 days later, elicitation was performed by applying 10 μL of 0.5% DNFB (1-fluoro-2,4-dinitrobenzene) in 1:4 olive oil:acetone to the right ear and the vehicle mixture to the left ear of each mouse, under isoflurane anesthesia. The treated areas were allowed to dry for 4 minutes before the mice were revived. Hindleg nails were clipped to prevent subsequent scratching of the elicited site (ear).

### Dex Treatment of DNFB-Elicited Mice

One hour prior to elicitation, mice were administered Dex by i.p. injection (2.8 mg/kg, with 3:7 PEG400:PBS as vehicle). 0.45% Dex was also combined with the 0.5% DNFB solution (1:4 olive oil:acetone) applied topically to induce elicitation. Control mice received i.p. injection of the vehicle alone and were treated topically with 0.5% DNFB solution in 1:4 olive oil:acetone.

### Evans Blue Assay

Isoflurane anesthetized mice were i.v. injected (retro-orbital) with 100 μL of 1% Evans Blue dye in PBS, filtered through 0.45 μm strainer. The mice were euthanized 1.5 hours later, at which point their ears were harvested and placed into 1 mL of formamide for 2 days at 56 °C. Evans Blue levels were then quantified as the difference between absorbance at 620 nm and 720 nm.

### Ear Thickness Measurements

Two days post challenge, DNFB-elicited mice were anaesthetized with isoflurane, and their ear thickness was measured using the Dial Thickness Gauge (Mitutoyo, Manufacturer Part Number: 7300A).

### Chemical Depilation

Ear skin was depilated 10-14 days prior to the initiation of any experimental protocol. Mice were first anesthetized with isoflurane. Mouse ears were coated with a thin layer of Nair^TM^ and rinsed with PBS following a 4-minute incubation. Ear skin was then dried using a kimwipe. Hind leg nails were clipped to prevent subsequent scratching of the chemically depilated site.

### Whole tissue RNA isolation and Quantitative RT-PCR

Ears were harvested from euthanized mice and mechanically homogenized in TRIzol using razor blades, followed by bead lysis. RNA was isolated from the TRIzol mixture in accordance with the TRIzol Reagent User guide (Doc. Part No.15596026.PPS, Pub. No.MAN0001271). The resulting RNA was then purified using the Zymo Research RNA Clean & Concentrator kit per the manufacturer’s instructions. RNA purity and concentration were determined using a NanoDrop spectrophotometer. cDNA was synthesized from total RNA and then used as a template for qRT-PCR using the LunaScript® Universal One Step Kit on a StepOnePlus device (Applied Biosystems). mRNA expression levels were normalized to *Gapdh* expression and relative gene expression between samples was calculated using the -ΔΔCt method. Primer sequences are listed in Table 1.

**Table 1.**
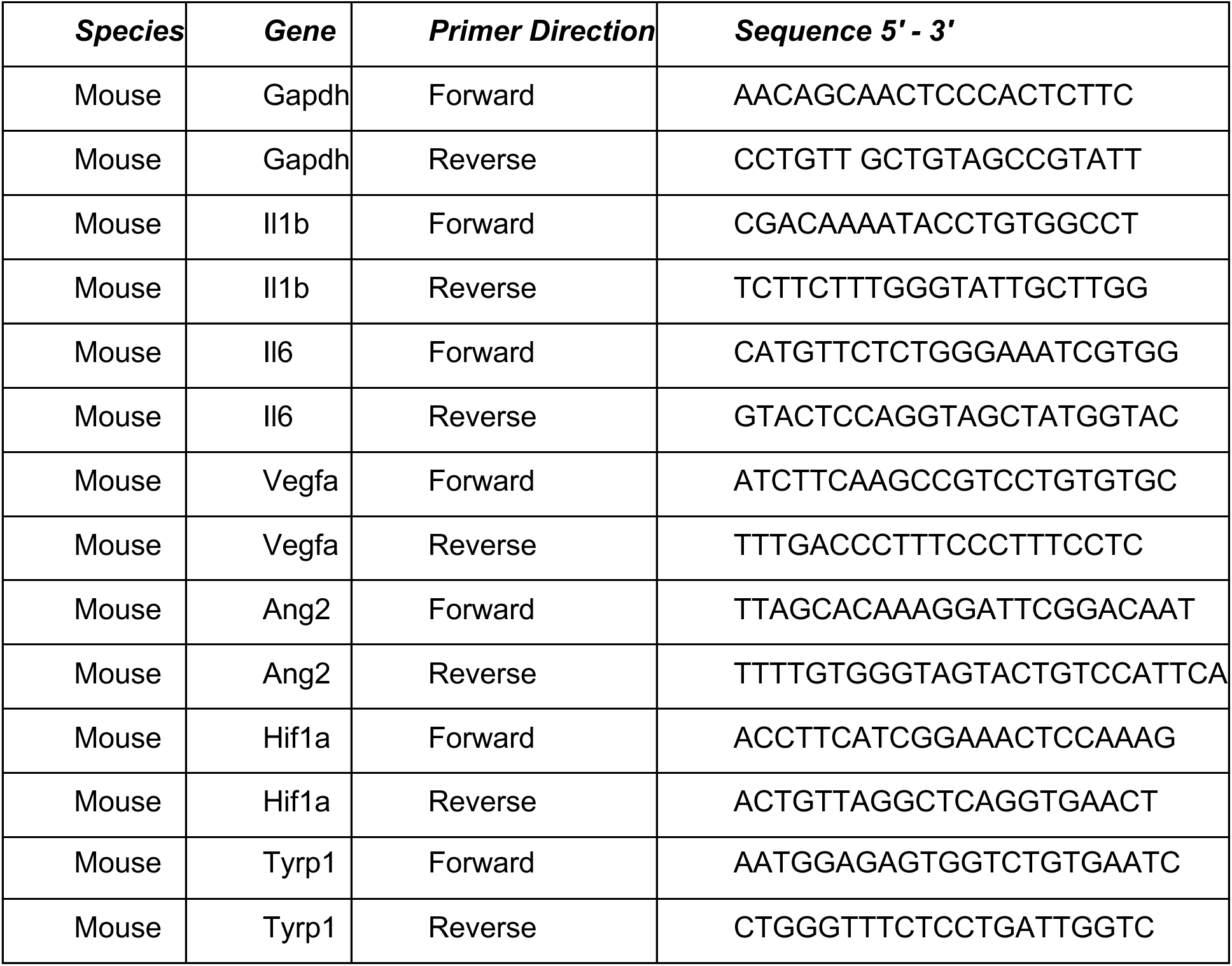
List of qRT-PCR Primers.

### Macroscopic Photography

A NikonZ6 camera fitted with Nikon 1.5 mm macro lens was used to capture all macroscopic images of mouse ears. Photographs were processed using Adobe Lightroom Classic (v14.1.1) and Microsoft PowerPoint. All compared images received identical processing.

### Generation of bone marrow (BM) chimeras

*BPT;LSL-TdTomato* mice were lethally irradiated (900 cGy) using a cesium source and reconstituted with 4 × 10^6^ bone marrow cells derived from *Cd11c-yfp* donor mice. Reconstitution of the hematopoietic compartment was monitored in the blood starting 6 weeks after reconstitution.

### Intravital Imaging of Murine Ears

For intravital imaging studies, mice were anaesthetized and maintained on 1.5% isoflurane in 100% oxygen for up to 2 hours. Each mouse was restrained on a stage warmer at 37°C (BioTherm Micro S37; Biogenics) and the ear was mounted onto a custom-made stage for imaging^75^. Two-photon microscopy was performed on an Olympus FVMPE- RS system equipped with a 25× 1.05 NA water immersion objective and a tunable InSight laser. Images were acquired with Fluoview software (FVS31). For experiments using *BPT-LSL-TdTomato::Cd11c-yfp* mice, YFP and RFP were detected with 1020 nm excitation using the Yellow/Red (515–565 nm) filter, and second harmonics (collagen) were detected with 1020 nm excitation using the Blue/Green (460–500 nm) filters, separated by an LP 520 dichroic mirror. Overlapping images were assembled using the same acquisition software. For blood vessel tracing experiments using *BPT-LSL-TdTomato* mice, 950 nm excitation was used. FITC and second harmonics (collagen) was detected using the Blue/Green (460–500 nm) filter, and TdTomato was detected using the Green/Red (495–500 nm) filter, separated by an LP 570 dichroic mirror. We typically acquired 150-200 μm-deep Z-stacks with 4 μm Z resolution, 12 tiles (4×3), with an X-Y resolution of 800 × 800 pixels per tile, and 0.6364 μm per pixel. Post-production processing of images was done using Fiji (v2.9.0), including tile stitching and quantification of GFP signal. FITC-Dextran Signal Area was calculated as the percentage of total GFP pixels above threshold in a maximum Z-projected image.

### Immune Cell Isolation

Lymph nodes were mechanically disrupted and filtered through a 70-μm strainer to create single cell suspensions in cold complete RPMI (cRPMI) (RPMI 1640 media supplemented with 2 μM glutamine, 100 U/ml penicillin/streptomycin, and 10% FBS). Ears were mechanically dissociated and then digested in 100 μg/mL Liberase, 500 μg/mL Hyaluronidase, 10 mM HEPES, in up to 5 mL serum-free DMEM, incubated for 1 hour at 37°C, and filtered over 70 μm nylon mesh to create single cell suspensions in cold cRPMI. Isolated B16F10 tumors were mechanically dissociated and then digested in 1 mg/mL Collagenase III for 1 hour at 37°C, and then filtered through a 70-μm strainer. Tumor homogenate was then centrifuged over a 40/80% Percoll gradient to isolate immune cells.

### Intracellular cytokine and transcription factor staining

Intracellular cytokine staining was performed using the Cytofix/Cytoperm PlusKit per manufacturer’s instructions. In brief, single-cell suspensions from freshly harvested dLN were stimulated with PMA (20 ng/mL) and ionomycin (1 μg/mL) for 4 hours at 37 °C in the presence of GolgiPlug (brefeldin A). After staining for cell-surface molecules, the cells were fixed, permeabilized, and stained with antibodies to IFNγ, IL-17a, and IL-4. Flow cytometry antibodies are listed in Table 2.

**Table 2.**
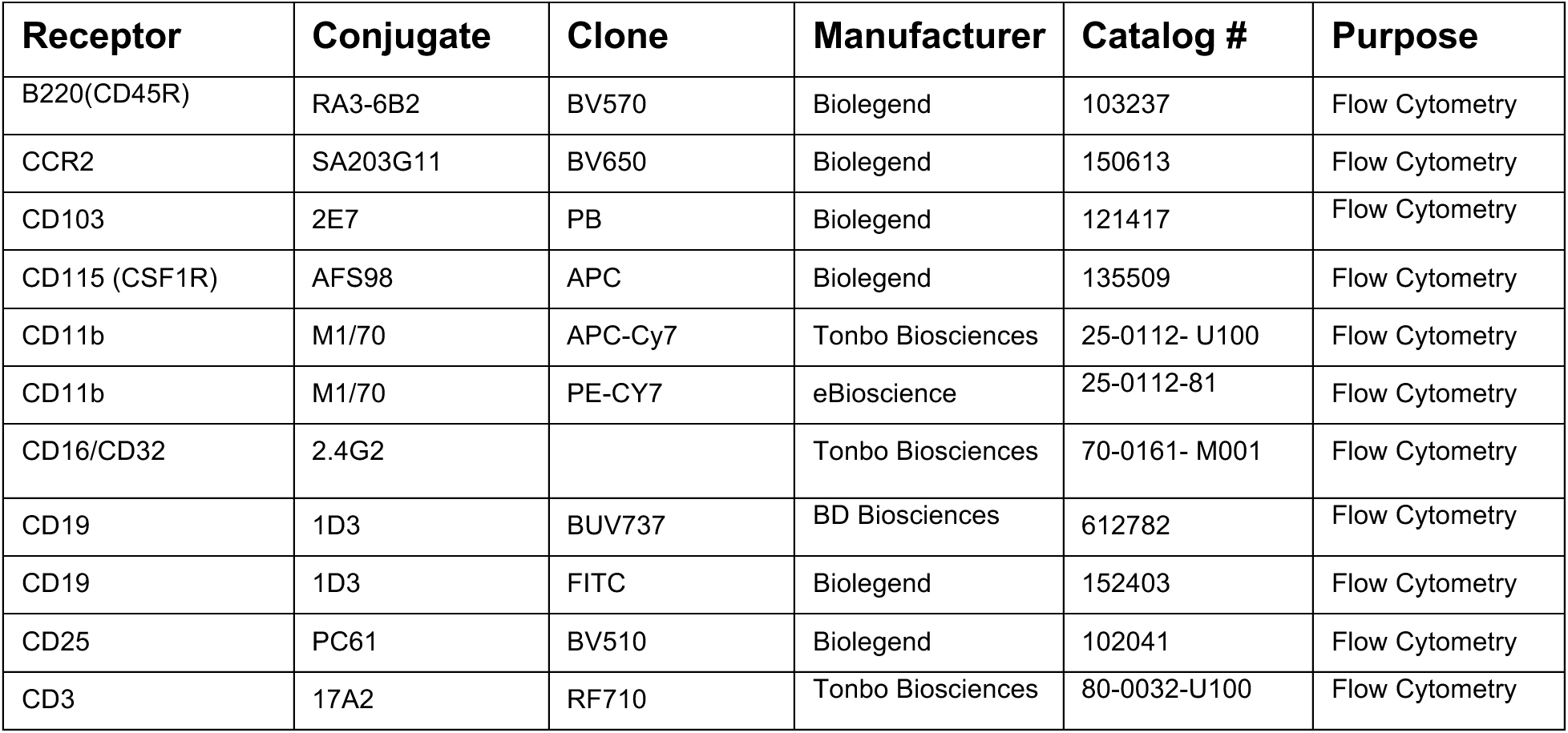
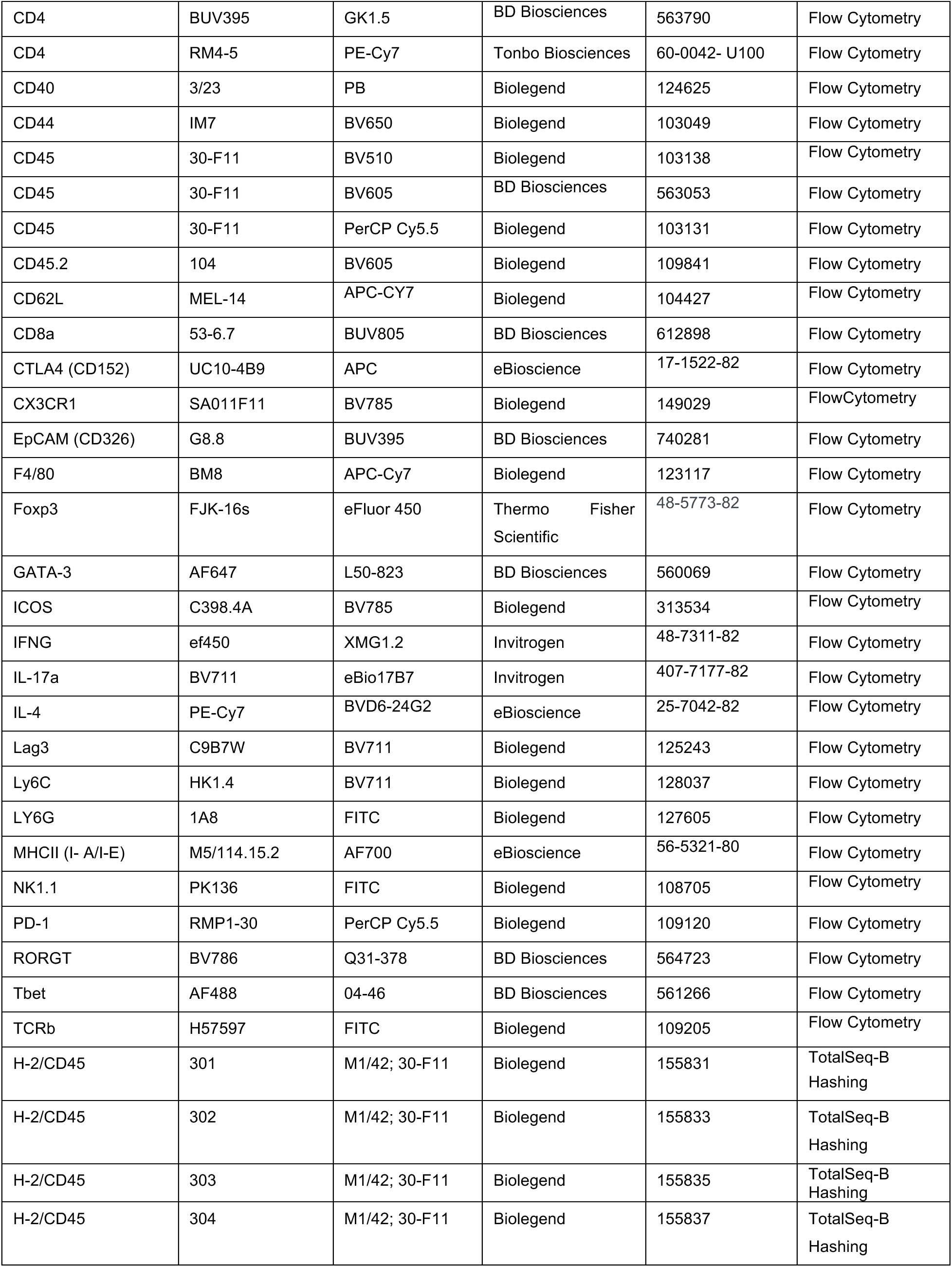
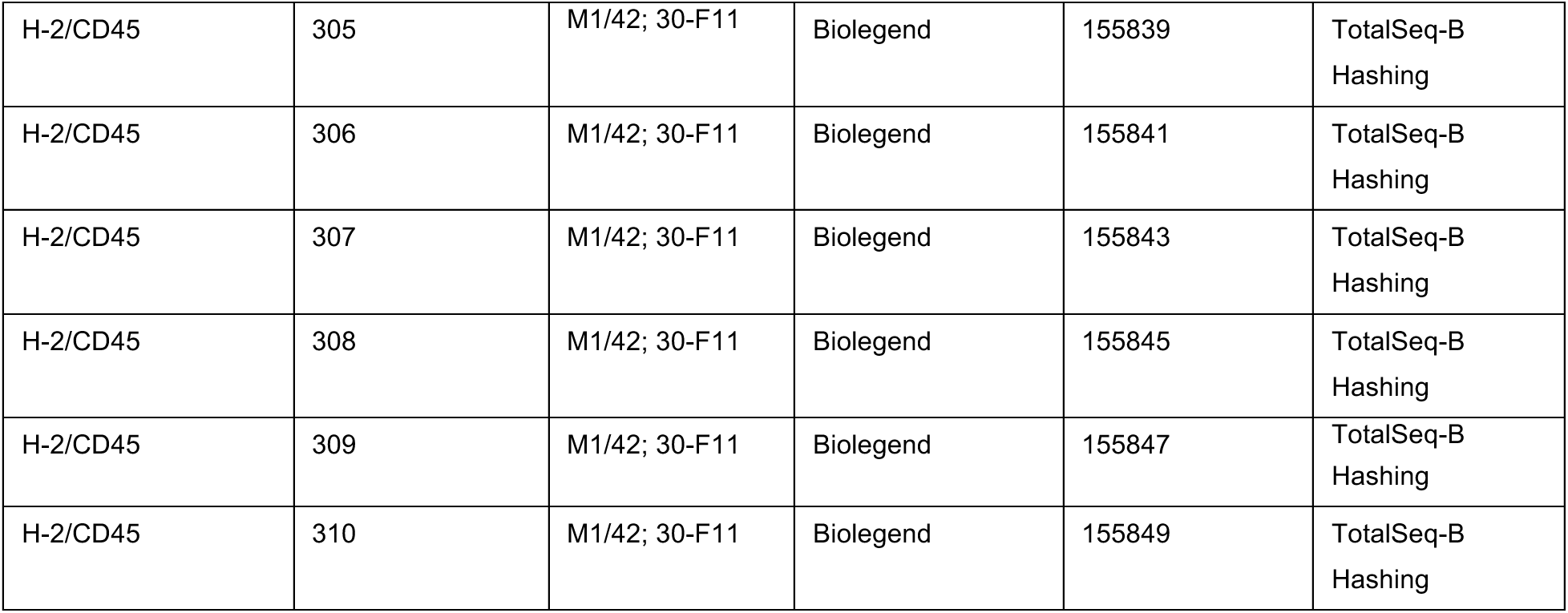
List of Antibodies.

### Flow cytometry

Single cell suspensions were washed and resuspended with FACS buffer (PBS, 2% FBS, 0.02% Sodium Azide) containing purified rat anti-mouse CD16/CD32 (Fc Block). Cells were then stained with antibody cocktails for 20 minutes at 4°C. When necessary, intracellular Foxp3 staining was performed using a Foxp3 mouse T_reg_ cell staining kit (eBioscience). Cells were washed with FACS buffer after staining, resuspended in FACS buffer, and analyzed using a Cytek Biosciences Aurora Flow Cytometer. Flow cytometry data was analyzed with FlowJo (v10.0) software (BD Biosciences).

### Quantification and statistical analysis

Unless otherwise stated, data in graphs are shown as mean ± standard deviation (SD). Statistical analyses were performed using Prism 10 (GraphPad v10.1.1). Statistical tests (e.g. one-way ANOVA, paired t-test, etc.), sample size (n) and what n represents (e.g. number of mice) are documented in the figure legends.

### Single-Cell RNA Sequencing

Single-cell RNA sequencing was performed on FACS-sorted cell populations using the Chromium instrument (10x Genomics) following the manufacturer’s protocol for 3′ v3 chemistry. Briefly, single-cell suspensions were washed and resuspended in FACS buffer containing antibodies against rat anti-mouse CD45 and/or rat anti-mouse Ly6G, followed by staining for 30 minutes at 4°C. Cells were then washed, resuspended in FACS buffer, and sorted using an Aria II cytometer (BD Biosciences). Sorted cells were subsequently stained with TotalSeqTM-B barcoded hashing antibodies for sample multiplexing (Table 2). Immune cells from each condition were pooled at equal ratios, washed twice with PBS containing 0.05% bovine serum albumin (BSA), and resuspended in PBS with 0.04% BSA to a final concentration of 700–1,300 cells per μL. Cell viability was confirmed to be above 70% using 0.2% (w/v) Trypan Blue staining (Countess II). Library preparation of single-cell suspensions containing hash-tagged cells was performed by the MSKCC Integrated Genomics Operation (IGO) Core Facility. Subsequent cell capture, reverse transcription, cDNA amplification, and library construction were carried out according to the standard 10x Genomics protocol. Raw sequencing data were processed using the 10x Genomics

Cell Ranger pipeline. The Cell Ranger mkfastq tool was used to demultiplex raw sequencing files (BCL format) into FASTQ files. Alignment of gene expression libraries to the GRCm39 reference transcriptome, generation of count matrices, and processing of feature barcode and VDJ library reads, were performed using the Cell Ranger (v9.0) default pipeline. For downstream analysis, the Seurat package (v5.2.0) in R (v4.2.3) was used. Data from all scRNA-seq samples were merged into a single aggregated Seurat object using IntegrateData()^76^. Hashtags were demultiplexed using HTODemux()^77^. The standard Seurat workflow was followed, including quality control, normalization, feature selection, dimensionality reduction, clustering, and differential expression analysis. Immune cell subsets were annotated against the Immgen database using the SingleR R package^78^. Cell lineage trajectories and pseudotime values were inferred using Slingshot^79^.

## Acknowledgements

We thank A. Y. Rudensky, J. Wolchok, and T. Merghoub for mice; A. Schietinger and M. O. Li for conceptual guidance; G. Diehl for critical reading of the manuscript; the MSKCC Flow Cytometry Core Facility for FACS purification; the MSKCC Integrated Genomics Operation for library preparation and sequencing; and members of the M. Huse lab for advice. Supported in part by the US National Institutes of Health (R01-AI087644 to M. H., R01-CA286566 to M. H., R01-AR077664 to J. H. Z., F31-CA294974 to Y. A. E., P30-CA008748 to MSKCC), the Ludwig Center for Cancer Immunotherapy (M. H.), the Vision of Children Foundation (J. H. Z.), and the National Organization for Albinism and Hypopigmentation (J. H. Z.).

---

**Figure 1- figure supplement 1.**
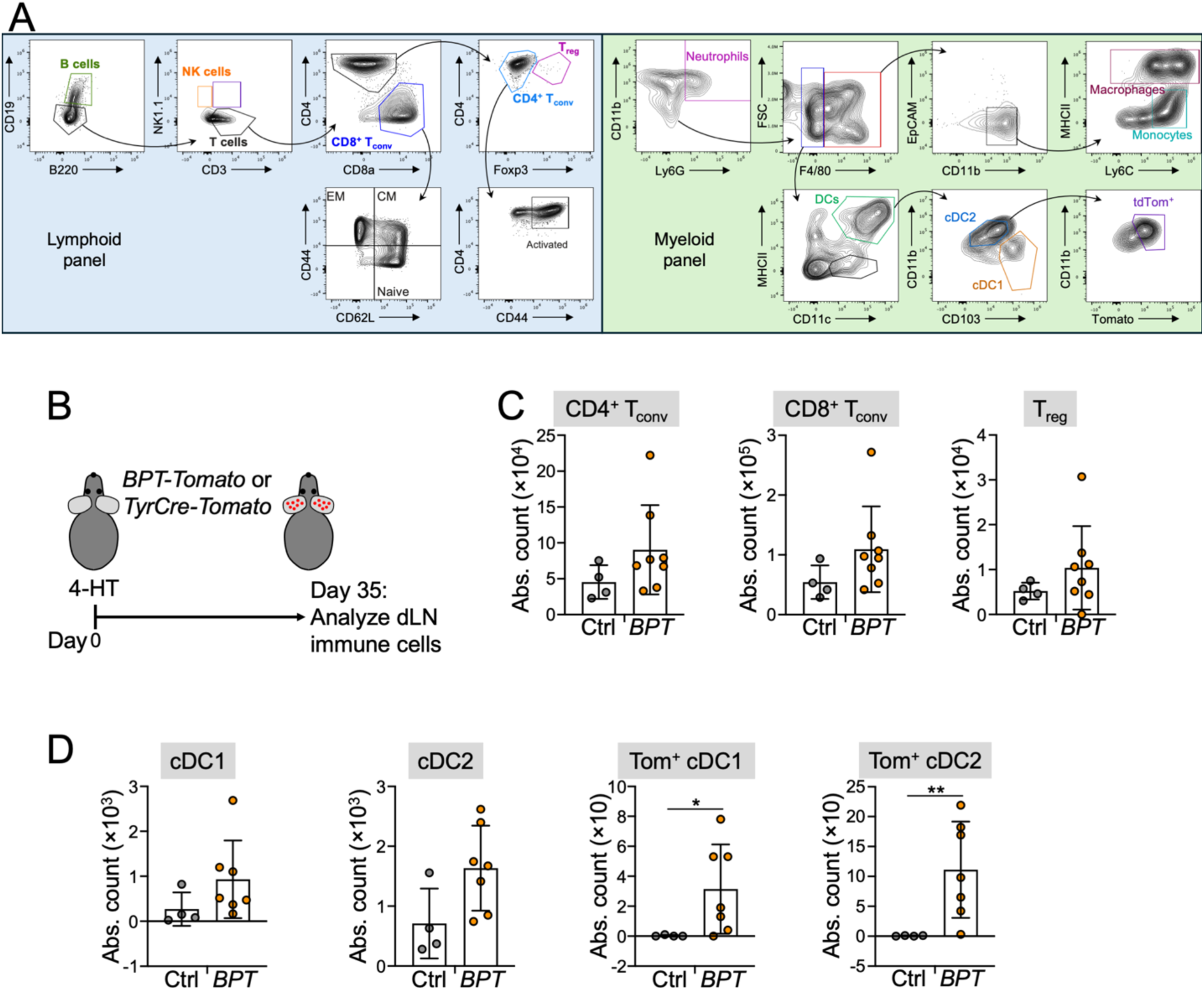
Flow cytometric analysis of *BPT* mice. (A) Gating strategies for lymphoid (left) and myeloid (right) panels, starting from live, CD45^+^ singlets. (B-C) *BPT-TdTomato* and *Tyr-TdTomato* control mice were 4-HT-painted and CD45^+^ cells in the dLN enumerated after 35 days by flow cytometry. (B) Schematic diagram of the experimental approach. (C) Quantification of the indicated T cell subsets. (D) Quantification of the indicated myeloid cell subsets. Tom^+^ = TdTomato positive. * and ** denote P ≤ 0.05 and P < 0.01, respectively, calculated by Mann-Whitney test.

**Figure 2 - supplement 1.**
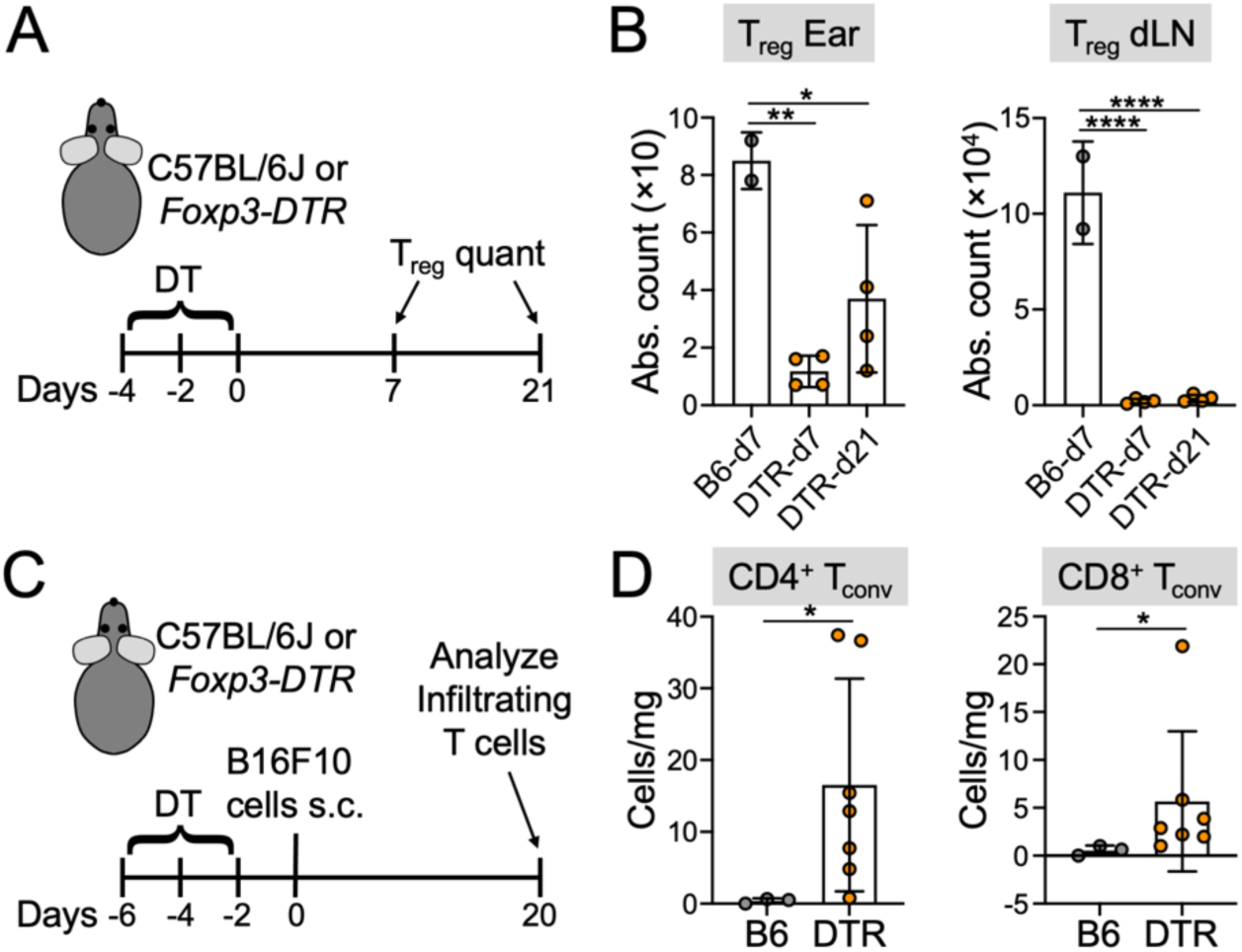
Flow cytometric analysis of T_reg_ cell depletion and T_conv_ cell infiltration in *Foxp3-DTR* mice. (A-B) *Foxp3-DTR* mice and C57BL/6J controls were DT-treated and T_reg_ cells quantified by flow cytometry in the ear skin and dLN 7 and 21 days after the last DT dose. (A) Schematic diagram of the experimental approach. (B) Absolute count of T_reg_ cells in the ear skin (left) and dLN (right). Error bars indicate SD. *, **, and **** denote P ≤ 0.05, P < 0.01, and P < 0.0001, respectively, calculated by one-way ANOVA. N = 2 C57BL/6J control mice and 4 *Foxp3-DTR* mice. (C-D) *Foxp3-DTR* mice and C57BL/6J controls were DT-treated and then injected s.c. with B16F10 cells. 20 days later, tumor infiltrating T cells were quantified by flow cytometry. (C) Schematic diagram of the experimental approach. (D) Quantification of tumor infiltrating CD4^+^ (left) and CD8^+^ (right) T_conv_ cells. Error bars indicate SD. * denotes P ≤ 0.05, calculated by lognormal Welch’s t-test. N = 3 C57BL/6J control mice and 7 *Foxp3-DTR* mice.

**Figure 3 - supplement 1.**
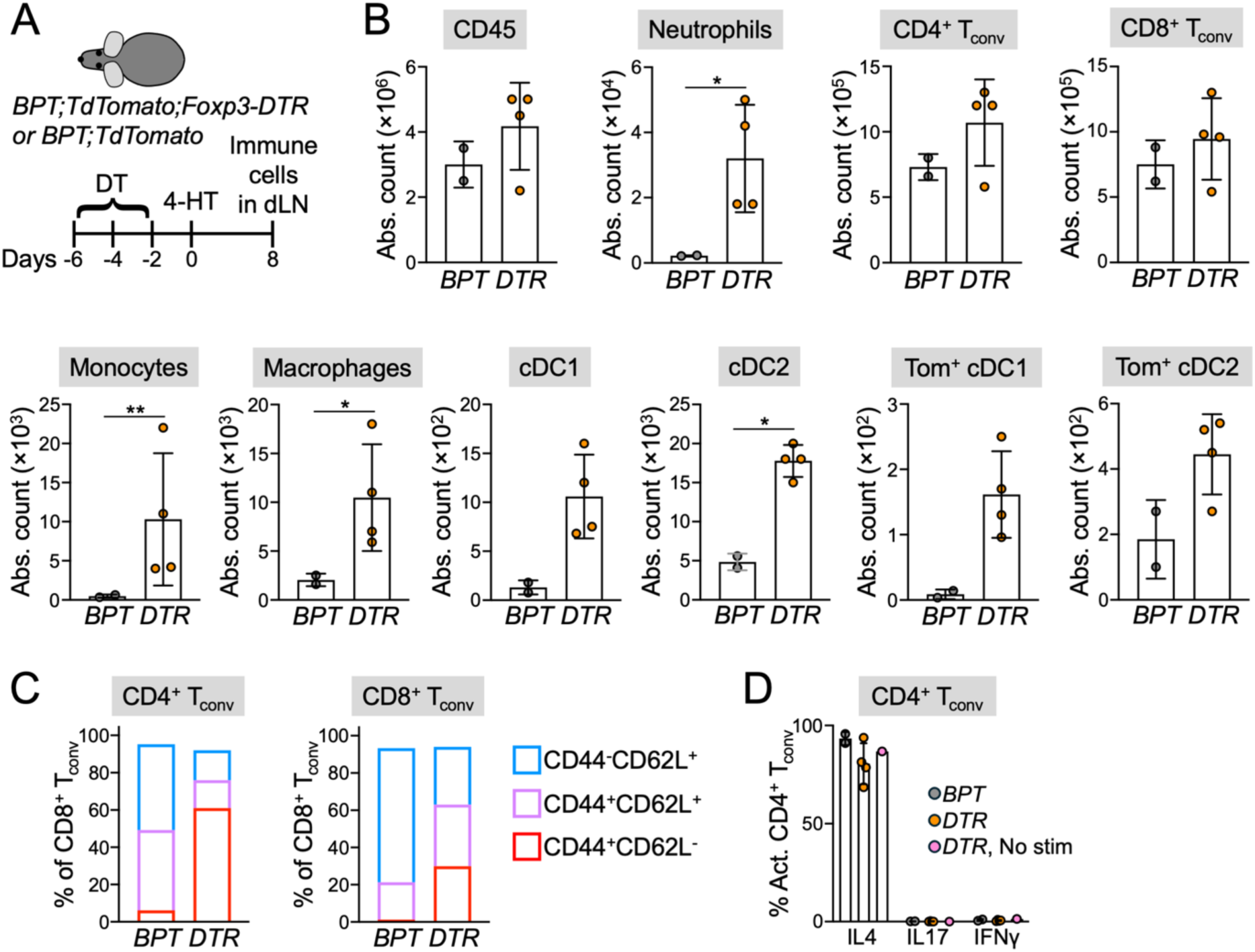
T_reg_ cell depletion drives multimodal inflammation in the dLN. *BPT-TdTomato;Foxp3-DTR* mice and *BPT-TdTomato* controls were treated with DT, followed by 4-HT painting, and immune cells in the dLN enumerated by flow cytometry after 8 days. (A) Schematic of the experimental protocol. (B) Quantification of the indicated immune cell subsets. Tom^+^ = TdTomato positive. * and ** denote P ≤ 0.05 and P < 0.01, respectively, calculated by lognormal Welch’s t-test. N = 2 *BPT-TdTomato* mice and 4 *BPT-TdTomato;Foxp3-DTR* mice. (C) Subset analysis of CD4^+^ (left) and CD8^+^ (right) T_conv_ cells in the dLN at the experimental endpoint. Fractions of total represent mean values derived from N = 2 *BPT* mice and N = 4 *BPT;Foxp3-DTR* mice. (D) Intracellular cytokine staining by CD4^+^ T_conv_ cells after *in vitro* stimulation with PMA/ionomycin. Error bars indicate SD. N = 2 *BPT* mice and N = 4 *BPT;Foxp3-DTR* mice.

**Figure 4 - supplement 1.**
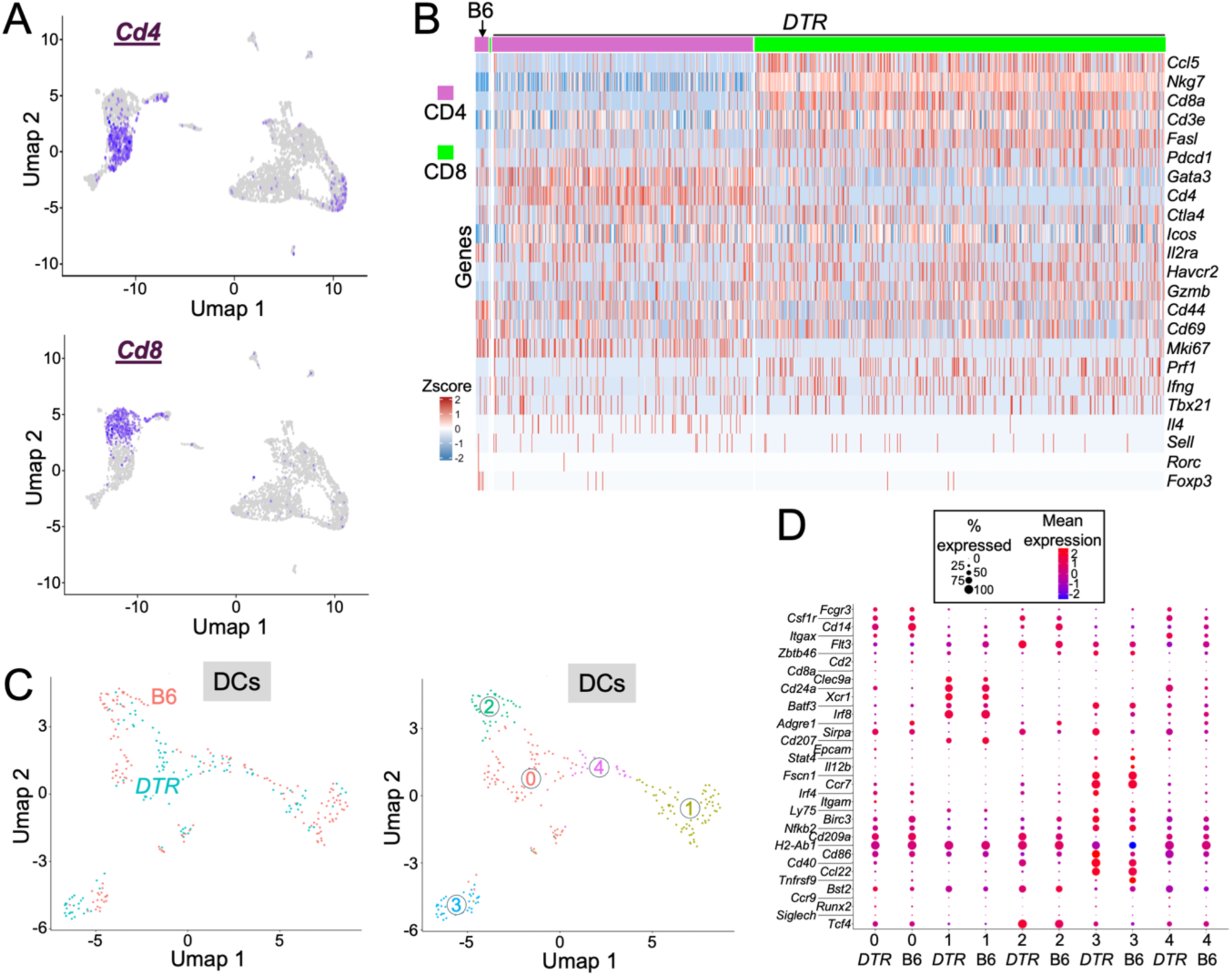
T_reg_ cell depletion drives multimodal inflammation in the skin. CD45^+^cells extracted from the ear skin of *Foxp3-DTR* mice or C57BL/6J controls were analyzed by scRNA-seq 8 days after DT treatment. (A) Feature plots of the all cell UMAP, showing expression of *Cd4* and *Cd8*. (B) Gene expression heat map of Immgen-defined T cells, organized by sample (CD57BL/6J (B6) or *Foxp3-DTR* (DTR)) and by lineage (CD4 or CD8). (C) UMAP reclustering of the DCs identified in the all cell UMAP. Cells are colored by sample on the left and by Seurat cluster on the right. (D) Dot plot showing the expression of critical DC lineage genes, organized by sample and Seurat cluster.

**Figure 4 - supplement 2.**
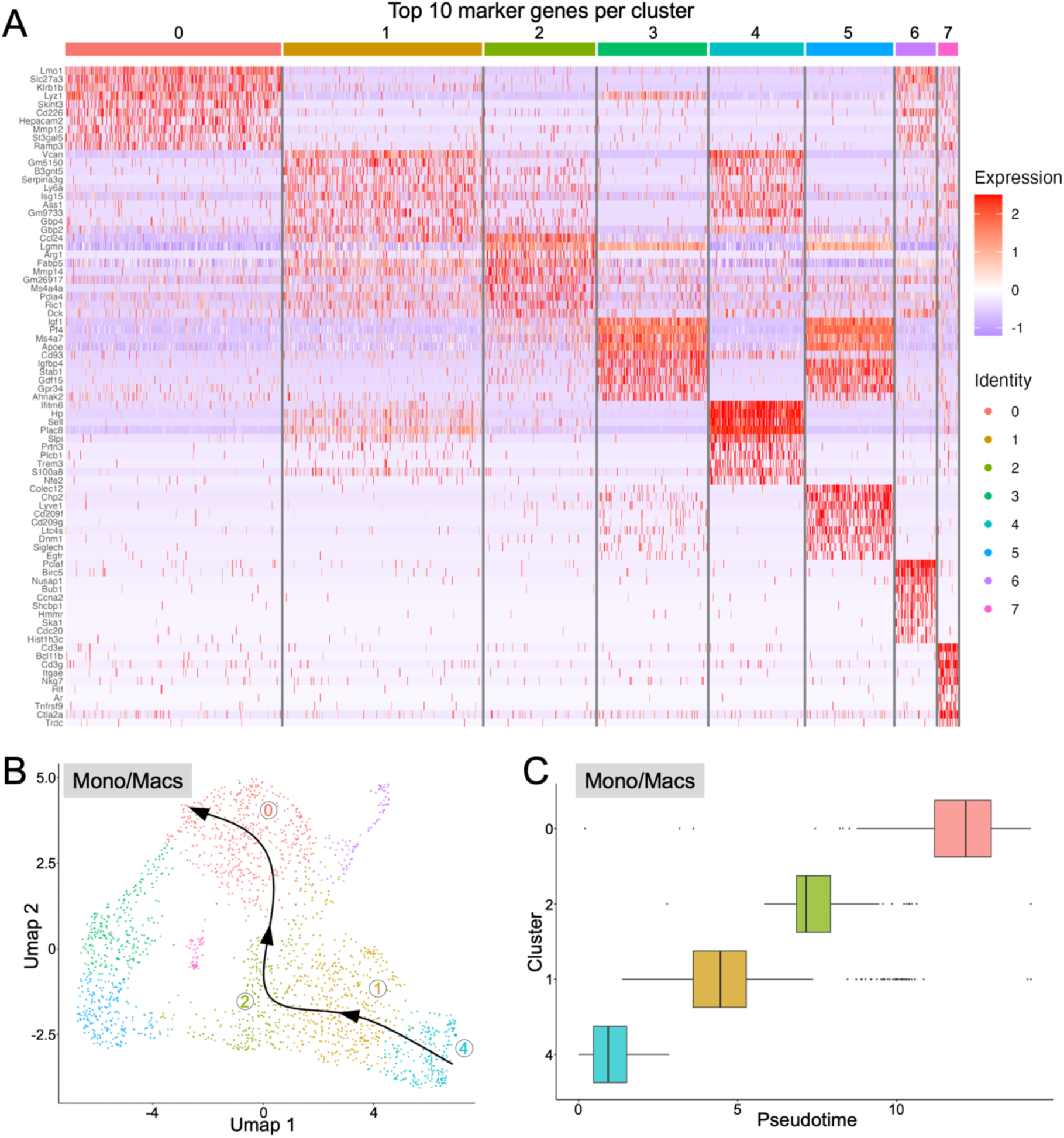
Inflammatory monocyte/macrophage recruitment and differentiation in T_reg_ cell-depleted skin. CD45^+^ cells extracted from the ear skin of *Foxp3-DTR* mice or C57BL/6J controls were analyzed by scRNA-seq 8 days after DT treatment. Monocyte and macrophage clusters from the all cell analysis were subjected to Seurat reclustering. (A) Heat map showing the differentially expressed genes that define the resulting monocyte/macrophage clusters. (B-C) Pseudotime analysis was performed on clusters 0, 1, 2, and 4. (B) Pseudotime trajectory output mapped onto the monocyte/macrophage UMAP. (C) The pseudotime distribution of each cluster, with central line denoting median, colored boxes denoting the central 50^th^ percentile, and whiskers indicating the range, with outliers (determined by Tukey’s method) shown as individual dots.

**Figure 6 - supplement 1.**
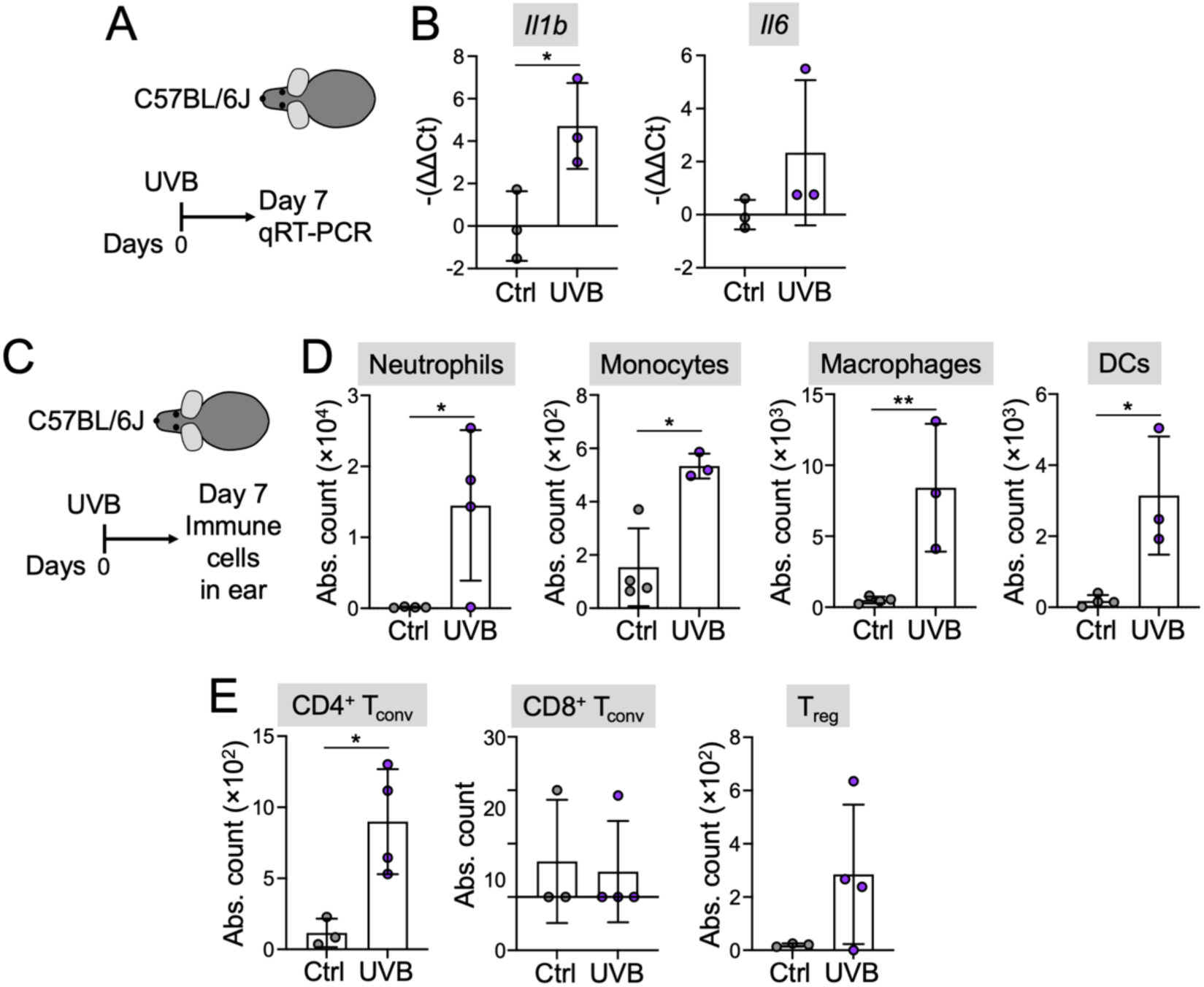
UVB induces cutaneous inflammation. (A-B) C57BL/6J mice were UVB- or mock-irradiated and qRT-PCR performed on skin homogenates 7 days later. (A) Schematic of the experimental protocol. (B) Quantified expression of the indicated genes. * denotes P ≤ 0.05, calculated by unpaired t-test. N ≥ 3 mice per group. (C-E) C57BL/6J mice were UVB- or mock-irradiated and immune cells in the ear skin enumerated by flow cytometry after 8 days. (C) Schematic of the experimental protocol. (D) Quantification of the indicated myeloid cell subsets. (E) Quantification of the indicated T cell subsets. In D and E, error bars indicate SD. * and ** denote P ≤ 0.05 and P < 0.01, respectively, calculated by lognormal Welch’s t-test. N ≥ 3 mice per group.

**Figure 6 - supplement 2.**
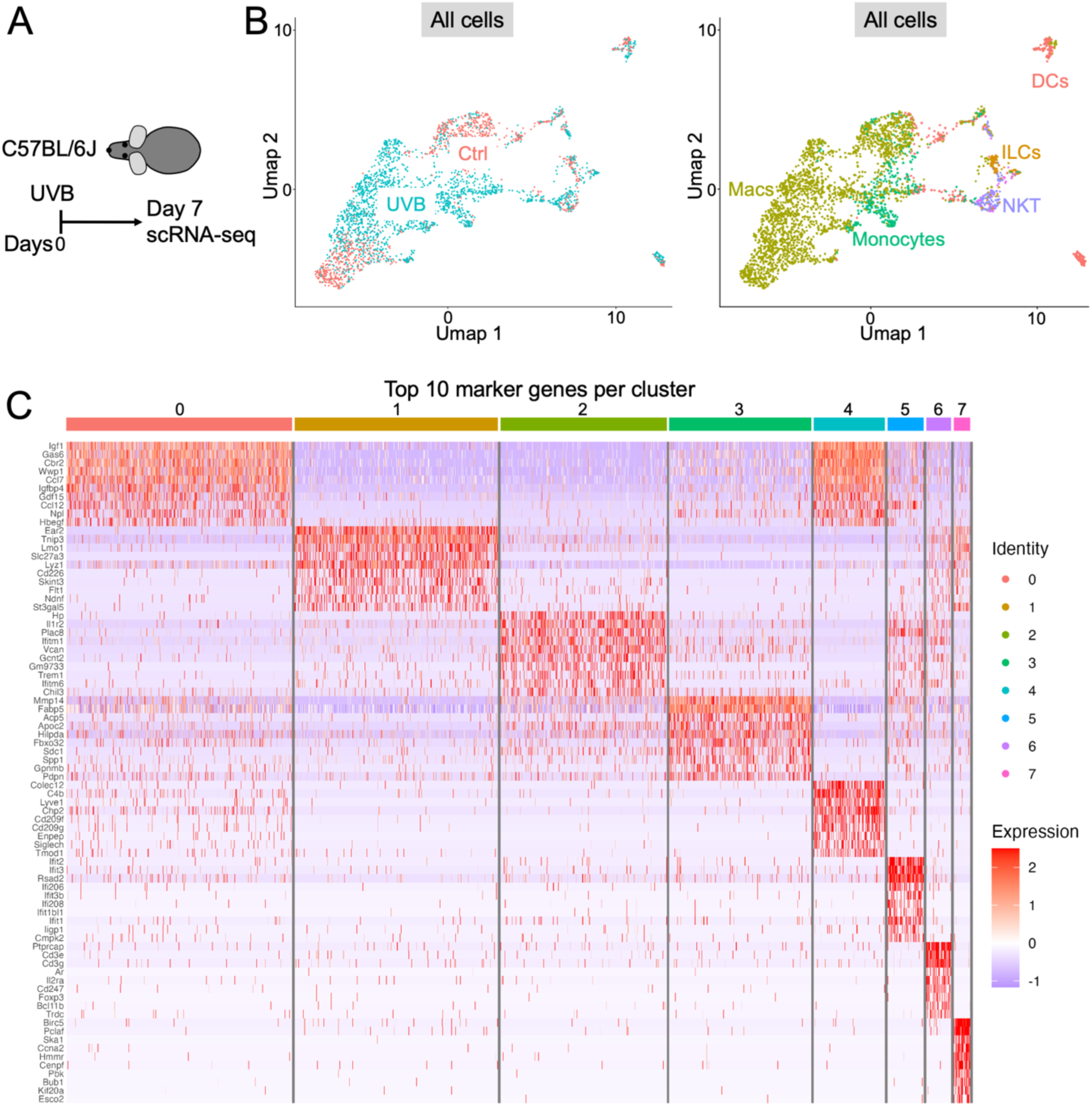
UVB irradiation drives multimodal inflammation in the skin. C57BL/6J mice were exposed to UVB or mock irradiation and CD45^+^ cells from the ear skin analyzed by scRNA-seq 7 days later. (A) Schematic of the experimental protocol. (B) UMAP showing all sequenced cells from both samples, colored by sample on the left and by cell type on the right. (C) Monocyte and macrophage clusters from the all cell analysis were subjected to Seurat reclustering. The heat map shows differentially expressed genes that define the resulting monocyte/macrophage clusters.

**Figure 7 - supplement 1.**
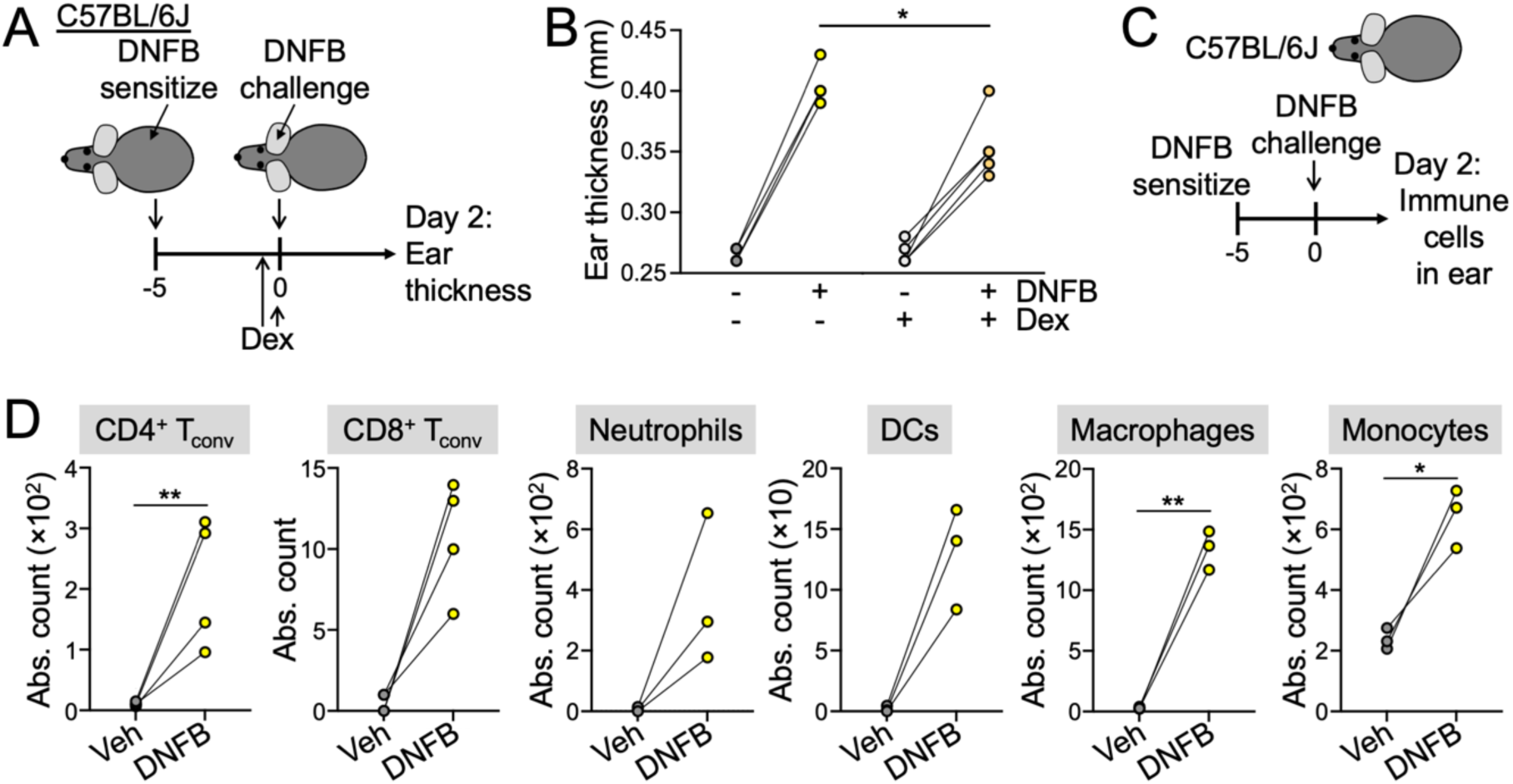
DNFB-induced hypersensitivity drives cutaneous inflammation. (A-B) C57BL/6J mice were sensitized with DNFB and then rechallenged on one ear in the presence or absence of Dex therapy. Ear thickness was measured 2 days after elicitation. (A) Schematic of the experimental protocol. (B) Ear thickness measurements, pairing DNFB-elicited ears with contralateral controls. N = 4 mice per group. * denotes P ≤ 0.05, calculated by unpaired t-test. (C-D) C57BL/6J mice were sensitized with DNFB and then rechallenged 5 days later. Immune cells in the ear were enumerated by flow cytometry 2 days after elicitation. (C) Schematic of the experimental protocol. (D) Infiltration of the indicated immune subsets, pairing DNFB-elicited ears with contralateral controls. N = 4 mice per group. * and ** denote P ≤ 0.05 and P < 0.01, respectively, calculated by paired lognormal t-test.

## References

1. Didier, A.J., Nandwani, S.V., Watkins, D., Fahoury, A.M., Campbell, A., Craig, D.J., Vijendra, D., and Parquet, N. (2024). Patterns and trends in melanoma mortality in the United States, 1999-2020. BMC Cancer 24, 790. 10.1186/s12885-024-12426-z.

2. Switzer, B., Puzanov, I., Skitzki, J.J., Hamad, L., and Ernstoff, M.S. (2022). Managing Metastatic Melanoma in 2022: A Clinical Review. JCO Oncol Pract 18, 335–351. 10.1200/OP.21.00686.

3. Leonardi, G.C., Falzone, L., Salemi, R., Zanghi, A., Spandidos, D.A., McCubrey, J.A., Candido, S., and Libra, M. (2018). Cutaneous melanoma: From pathogenesis to therapy (Review). Int J Oncol 52, 1071–1080. 10.3892/ijo.2018.4287.

4. Maverakis, E., Cornelius, L.A., Bowen, G.M., Phan, T., Patel, F.B., Fitzmaurice, S., He, Y., Burrall, B., Duong, C., Kloxin, A.M., et al. (2015). Metastatic melanoma - a review of current and future treatment options. Acta Derm Venereol 95, 516–524. 10.2340/00015555-2035.

5. Pollock, P.M., Harper, U.L., Hansen, K.S., Yudt, L.M., Stark, M., Robbins, C.M., Moses, T.Y., Hostetter, G., Wagner, U., Kakareka, J., et al. (2003). High frequency of BRAF mutations in nevi. Nat Genet 33, 19–20. 10.1038/ng1054.

6. Hayward, N.K., Wilmott, J.S., Waddell, N., Johansson, P.A., Field, M.A., Nones, K., Patch, A.M., Kakavand, H., Alexandrov, L.B., Burke, H., et al. (2017). Whole-genome landscapes of major melanoma subtypes. Nature 545, 175–180. 10.1038/nature22071.

7. Lozada, J.R., Geyer, F.C., Selenica, P., Brown, D., Alemar, B., Merghoub, T., Berger, M.F., Busam, K.J., Halpern, A.C., Weigelt, B., et al. (2019). Massively parallel sequencing analysis of benign melanocytic naevi. Histopathology 75, 29–38. 10.1111/his.13843.

8. Gil-Barrachina, M., Hernando, B., Perez-Pastor, G., Alegre-de-Miquel, V., Valenzuela-Onate, C., Minguez-Lujan, S., Monfort-Lanzas, P., Tomas-Bort, E., Marques-Torrejon, M.A., and Martinez-Cadenas, C. (2026). Genetic Evolution of Melanoma: Comparative Analysis of Candidate Gene Mutations in Healthy Skin, Nevi, and Tumors from the Same Patients. Int J Mol Sci 27. 10.3390/ijms27010532.

9. Lee, K.J., Kao, Y.C., Jiang, A., Prokofyeva, A., Tom, L.N., Jagirdar, K., Smit, D.J., Oey, H.M., Tan, J.M., Ainger, S.A., et al. (2026). Subclinical fields of BRAF V600E-mutant melanocytes populate human skin and are enriched around melanoma and naevi. Br J Dermatol 194, 301–310. 10.1093/bjd/ljaf412.

10. Maher, N.G., Scolyer, R.A., and Colebatch, A.J. (2023). Biology and genetics of acquired and congenital melanocytic naevi. Pathology 55, 169–177. 10.1016/j.pathol.2022.12.344.

11. Schreiber, R.D., Old, L.J., and Smyth, M.J. (2011). Cancer immunoediting: integrating immunity’s roles in cancer suppression and promotion. Science 331, 1565–1570. 10.1126/science.1203486.

12. Shalapour, S., and Karin, M. (2015). Immunity, inflammation, and cancer: an eternal fight between good and evil. J Clin Invest 125, 3347–3355. 10.1172/JCI80007.

13. Postow, M.A., Callahan, M.K., and Wolchok, J.D. (2015). Immune Checkpoint Blockade in Cancer Therapy. J Clin Oncol 33, 1974–1982. 10.1200/JCO.2014.59.4358.

14. Wei, S.C., Duffy, C.R., and Allison, J.P. (2018). Fundamental Mechanisms of Immune Checkpoint Blockade Therapy. Cancer Discov 8, 1069–1086. 10.1158/2159-8290.CD-18-0367.

15. Josefowicz, S.Z., Lu, L.F., and Rudensky, A.Y. (2012). Regulatory T cells: mechanisms of differentiation and function. Annu Rev Immunol 30, 531–564. 10.1146/annurev.immunol.25.022106.141623.

16. Bos, P.D., Plitas, G., Rudra, D., Lee, S.Y., and Rudensky, A.Y. (2013). Transient regulatory T cell ablation deters oncogene-driven breast cancer and enhances radiotherapy. J Exp Med 210, 2435–2466. 10.1084/jem.20130762.

17. Shabaneh, T.B., Molodtsov, A.K., Steinberg, S.M., Zhang, P., Torres, G.M., Mohamed, G.A., Boni, A., Curiel, T.J., Angeles, C.V., and Turk, M.J. (2018). Oncogenic BRAF(V600E) Governs Regulatory T-cell Recruitment during Melanoma Tumorigenesis. Cancer Res 78, 5038–5049. 10.1158/0008-5472.CAN-18-0365.

18. Binnewies, M., Mujal, A.M., Pollack, J.L., Combes, A.J., Hardison, E.A., Barry, K.C., Tsui, J., Ruhland, M.K., Kersten, K., Abushawish, M.A., et al. (2019). Unleashing Type-2 Dendritic Cells to Drive Protective Antitumor CD4(+) T Cell Immunity. Cell 177, 556–571 e516. 10.1016/j.cell.2019.02.005.

19. Klages, K., Mayer, C.T., Lahl, K., Loddenkemper, C., Teng, M.W.L., Ngiow, S.F., Smyth, M.J., Hamann, A., Huehn, J., and Sparwasser, T. (2010). Selective Depletion of Foxp3(+) Regulatory T Cells Improves Effective Therapeutic Vaccination against Established Melanoma. Cancer Research 70, 7788–7799. 10.1158/0008-5472.Can-10-1736.

20. Hanahan, D. (2026). Hallmarks of cancer-Then and now, and beyond. Cell 189, 2254–2277. 10.1016/j.cell.2025.12.049.

21. Ortega-Gomez, A., Perretti, M., and Soehnlein, O. (2013). Resolution of inflammation: an integrated view. EMBO Mol Med 5, 661–674. 10.1002/emmm.201202382.

22. Rivera, L.B., and Bergers, G. (2015). Intertwined regulation of angiogenesis and immunity by myeloid cells. Trends Immunol 36, 240–249. 10.1016/j.it.2015.02.005.

23. De Palma, M., Biziato, D., and Petrova, T.V. (2017). Microenvironmental regulation of tumour angiogenesis. Nat Rev Cancer 17, 457–474. 10.1038/nrc.2017.51.

24. Bernard, J.J., Gallo, R.L., and Krutmann, J. (2019). Photoimmunology: how ultraviolet radiation affects the immune system. Nat Rev Immunol 19, 688–701. 10.1038/s41577-019-0185-9.

25. Schwarz, T. (2010). The dark and the sunny sides of UVR-induced immunosuppression: photoimmunology revisited. J Invest Dermatol 130, 49–54. 10.1038/jid.2009.217.

26. Clydesdale, G.J., Dandie, G.W., and Muller, H.K. (2001). Ultraviolet light induced injury: immunological and inflammatory effects. Immunol Cell Biol 79, 547–568. 10.1046/j.1440-1711.2001.01047.x.

27. Alexandrov, L.B., Nik-Zainal, S., Wedge, D.C., Aparicio, S.A., Behjati, S., Biankin, A.V., Bignell, G.R., Bolli, N., Borg, A., Borresen-Dale, A.L., et al. (2013). Signatures of mutational processes in human cancer. Nature 500, 415–421. 10.1038/nature12477.

28. Viros, A., Sanchez-Laorden, B., Pedersen, M., Furney, S.J., Rae, J., Hogan, K., Ejiama, S., Girotti, M.R., Cook, M., Dhomen, N., and Marais, R. (2014). Ultraviolet radiation accelerates BRAF-driven melanomagenesis by targeting TP53. Nature 511, 478–482. 10.1038/nature13298.

29. Hodis, E., Watson, I.R., Kryukov, G.V., Arold, S.T., Imielinski, M., Theurillat, J.P., Nickerson, E., Auclair, D., Li, L., Place, C., et al. (2012). A landscape of driver mutations in melanoma. Cell 150, 251–263. 10.1016/j.cell.2012.06.024.

30. Shain, A.H., Joseph, N.M., Yu, R., Benhamida, J., Liu, S., Prow, T., Ruben, B., North, J., Pincus, L., Yeh, I., et al. (2018). Genomic and Transcriptomic Analysis Reveals Incremental Disruption of Key Signaling Pathways during Melanoma Evolution. Cancer Cell 34, 45–55 e44. 10.1016/j.ccell.2018.06.005.

31. Muller, H.K., Malley, R.C., McGee, H.M., Scott, D.K., Wozniak, T., and Woods, G.M. (2008). Effect of UV radiation on the neonatal skin immune system- implications for melanoma. Photochem Photobiol 84, 47–54. 10.1111/j.1751-1097.2007.00246.x.

32. Mo, X., Preston, S., and Zaidi, M.R. (2019). Macroenvironment-gene-microenvironment interactions in ultraviolet radiation-induced melanomagenesis. Adv Cancer Res 144, 1–54. 10.1016/bs.acr.2019.03.008.

33. Sample, A., and He, Y.Y. (2018). Mechanisms and prevention of UV-induced melanoma. Photodermatol Photoimmunol Photomed 34, 13–24. 10.1111/phpp.12329.

34. Gieniusz, E., Skrzydlewska, E., and Luczaj, W. (2024). Current Insights into the Role of UV Radiation-Induced Oxidative Stress in Melanoma Pathogenesis. Int J Mol Sci 25. 10.3390/ijms252111651.

35. Enomoto, A., Yoshihisa, Y., Yamakoshi, T., Ur Rehman, M., Norisugi, O., Hara, H., Matsunaga, K., Makino, T., Nishihira, J., and Shimizu, T. (2011). UV-B radiation induces macrophage migration inhibitory factor-mediated melanogenesis through activation of protease-activated receptor-2 and stem cell factor in keratinocytes. Am J Pathol 178, 679–687. 10.1016/j.ajpath.2010.10.021.

36. Mukhopadhyay, P., Ferguson, B., Muller, H.K., Handoko, H.Y., and Walker, G.J. (2016). Murine melanomas accelerated by a single UVR exposure carry photoproduct footprints but lack UV signature C>T mutations in critical genes. Oncogene 35, 3342–3350. 10.1038/onc.2015.386.

37. Bowman, R.L., Hennessey, R.C., Weiss, T.J., Tallman, D.A., Crawford, E.R., Murphy, B.M., Webb, A., Zhang, S., La Perle, K.M., Burd, C.J., et al. (2021). UVB mutagenesis differs in Nras- and Braf-mutant mouse models of melanoma. Life Sci Alliance 4. 10.26508/lsa.202101135.

38. Gandini, S., Sera, F., Cattaruzza, M.S., Pasquini, P., Picconi, O., Boyle, P., and Melchi, C.F. (2005). Meta-analysis of risk factors for cutaneous melanoma: II. Sun exposure. Eur J Cancer 41, 45–60. 10.1016/j.ejca.2004.10.016.

39. Weiss, T.J., Crawford, E.R., Posada, V., Rahman, H., Liu, T., Murphy, B.M., Arnold, T.E., Gray, S., Hu, Z., Hennessey, R.C., et al. (2023). Cell-intrinsic melanin fails to protect melanocytes from ultraviolet-mutagenesis in the absence of epidermal melanin. Pigment Cell Melanoma Res 36, 6–18. 10.1111/pcmr.13070.

40. Hennessey, R.C., Holderbaum, A.M., Bonilla, A., Delaney, C., Gillahan, J.E., Tober, K.L., Oberyszyn, T.M., Zippin, J.H., and Burd, C.E. (2017). Ultraviolet radiation accelerates NRas-mutant melanomagenesis: A cooperative effect blocked by sunscreen. Pigment Cell Melanoma Res 30, 477–487. 10.1111/pcmr.12601.

41. An, H.T., Yoo, J.Y., Lee, M.K., Shin, M.H., Rhie, G.E., Seo, J.Y., Chung, J.H., Eun, H.C., and Cho, K.H. (2001). Single dose radiation is more effective for the UV-induced activation and proliferation of melanocytes than fractionated dose radiation. Photodermatol Photoimmunol Photomed 17, 266–271. 10.1034/j.1600-0781.2001.170604.x.

42. Yamazaki, S., Nishioka, A., Kasuya, S., Ohkura, N., Hemmi, H., Kaisho, T., Taguchi, O., Sakaguchi, S., and Morita, A. (2014). Homeostasis of thymus-derived Foxp3+ regulatory T cells is controlled by ultraviolet B exposure in the skin. J Immunol 193, 5488–5497. 10.4049/jimmunol.1400985.

43. Yamazaki, S., Odanaka, M., Nishioka, A., Kasuya, S., Shime, H., Hemmi, H., Imai, M., Riethmacher, D., Kaisho, T., Ohkura, N., et al. (2018). Ultraviolet B-Induced Maturation of CD11b-Type Langerin(-) Dendritic Cells Controls the Expansion of Foxp3(+) Regulatory T Cells in the Skin. J Immunol 200, 119–129. 10.4049/jimmunol.1701056.

44. Hesterberg, R.S., Amorrortu, R.P., Zhao, Y., Hampras, S., Akuffo, A.A., Fenske, N., Cherpelis, B., Balliu, J., Vijayan, L., Epling-Burnette, P.K., and Rollison, D.E. (2018). T Regulatory Cell Subpopulations Associated with Recent Ultraviolet Radiation Exposure in a Skin Cancer Screening Cohort. J Immunol 201, 3269–3281. 10.4049/jimmunol.1800940.

45. Dankort, D., Curley, D.P., Cartlidge, R.A., Nelson, B., Karnezis, A.N., Damsky, W.E., You, M.J., DePinho, R.A., McMahon, M., and Bosenberg, M. (2009). Braf(V600E) cooperates with Pten loss to induce metastatic melanoma. Nature Genetics 41, 544–552. 10.1038/ng.356.

46. Roberts, E.W., Broz, M.L., Binnewies, M., Headley, M.B., Nelson, A.E., Wolf, D.M., Kaisho, T., Bogunovic, D., Bhardwaj, N., and Krummel, M.F. (2016). Critical Role for CD103(+)/CD141(+) Dendritic Cells Bearing CCR7 for Tumor Antigen Trafficking and Priming of T Cell Immunity in Melanoma. Cancer Cell 30, 324–336. 10.1016/j.ccell.2016.06.003.

47. Kim, J.M., Rasmussen, J.P., and Rudensky, A.Y. (2007). Regulatory T cells prevent catastrophic autoimmunity throughout the lifespan of mice. Nat Immunol 8, 191–197. 10.1038/ni1428.

48. Boothby, I.C., Kinet, M.J., Boda, D.P., Kwan, E.Y., Clancy, S., Cohen, J.N., Habrylo, I., Lowe, M.M., Pauli, M., Yates, A.E., et al. (2021). Early-life inflammation primes a T helper 2 cell-fibroblast niche in skin. Nature 599, 667–672. 10.1038/s41586-021-04044-7.

49. Bieber, K., Sun, S., Witte, M., Kasprick, A., Beltsiou, F., Behnen, M., Laskay, T., Schulze, F.S., Pipi, E., Reichhelm, N., et al. (2017). Regulatory T Cells Suppress Inflammation and Blistering in Pemphigoid Diseases. Front Immunol 8, 1628. 10.3389/fimmu.2017.01628.

50. Sontheimer, C., Liggitt, D., and Elkon, K.B. (2017). Ultraviolet B Irradiation Causes Stimulator of Interferon Genes-Dependent Production of Protective Type I Interferon in Mouse Skin by Recruited Inflammatory Monocytes. Arthritis Rheumatol 69, 826–836. 10.1002/art.39987.

51. Toichi, E., Lu, K.Q., Swick, A.R., McCormick, T.S., and Cooper, K.D. (2008). Skin-infiltrating monocytes/macrophages migrate to draining lymph nodes and produce IL-10 after contact sensitizer exposure to UV-irradiated skin. J Invest Dermatol 128, 2705–2715. 10.1038/jid.2008.137.

52. Zhang, A.Y., Wu, C., Zhou, L., Ismail, S.A., Tao, J., McCormick, L.L., Cooper, K.D., and Gilliam, A.C. (2006). Transduced monocyte/macrophages targeted to murine skin by UV light. Exp Dermatol 15, 51–57. 10.1111/j.0906-6705.2005.00394.x.

53. Bernard, J.J., Cowing-Zitron, C., Nakatsuji, T., Muehleisen, B., Muto, J., Borkowski, A.W., Martinez, L., Greidinger, E.L., Yu, B.D., and Gallo, R.L. (2012). Ultraviolet radiation damages self noncoding RNA and is detected by TLR3. Nat Med 18, 1286–1290. 10.1038/nm.2861.

54. Schwarz, T., and Luger, T.A. (1989). Effect of UV irradiation on epidermal cell cytokine production. J Photochem Photobiol B 4, 1–13. 10.1016/1011-1344(89)80097-1.

55. Yano, K., Kadoya, K., Kajiya, K., Hong, Y.K., and Detmar, M. (2005). Ultraviolet B irradiation of human skin induces an angiogenic switch that is mediated by upregulation of vascular endothelial growth factor and by downregulation of thrombospondin-1. Br J Dermatol 152, 115–121. 10.1111/j.1365-2133.2005.06368.x.

56. Kim, M.S., Kim, Y.K., Eun, H.C., Cho, K.H., and Chung, J.H. (2006). All-trans retinoic acid antagonizes UV-induced VEGF production and angiogenesis via the inhibition of ERK activation in human skin keratinocytes. J Invest Dermatol 126, 2697–2706. 10.1038/sj.jid.5700463.

57. Di Nuzzo, S., Sylva-Steenland, R.M., de Rie, M.A., Das, P.K., Bos, J.D., and Teunissen, M.B. (1998). UVB radiation preferentially induces recruitment of memory CD4+ T cells in normal human skin: long-term effect after a single exposure. J Invest Dermatol 110, 978–981. 10.1046/j.1523-1747.1998.00220.x.

58. Hatton, J.L., Parent, A., Tober, K.L., Hoppes, T., Wulff, B.C., Duncan, F.J., Kusewitt, D.F., VanBuskirk, A.M., and Oberyszyn, T.M. (2007). Depletion of CD4+ cells exacerbates the cutaneous response to acute and chronic UVB exposure. J Invest Dermatol 127, 1507–1515. 10.1038/sj.jid.5700746.

59. Schwarz, A., Philippsen, R., and Schwarz, T. (2023). Mouse Models of Allergic Contact Dermatitis: Practical Aspects. J Invest Dermatol 143, 888–892. 10.1016/j.jid.2023.03.1668.

60. Fyhrquist, N., Lehtimaki, S., Lahl, K., Savinko, T., Lappetelainen, A.M., Sparwasser, T., Wolff, H., Lauerma, A., and Alenius, H. (2012). Foxp3+ cells control Th2 responses in a murine model of atopic dermatitis. J Invest Dermatol 132, 1672–1680. 10.1038/jid.2012.40.

61. Yamaguchi, H.L., Yamaguchi, Y., and Peeva, E. (2023). Role of Innate Immunity in Allergic Contact Dermatitis: An Update. Int J Mol Sci 24. 10.3390/ijms241612975.

62. Lugano, R., Ramachandran, M., and Dimberg, A. (2020). Tumor angiogenesis: causes, consequences, challenges and opportunities. Cell Mol Life Sci 77, 1745–1770. 10.1007/s00018-019-03351-7.

63. Togashi, Y., Shitara, K., and Nishikawa, H. (2019). Regulatory T cells in cancer immunosuppression - implications for anticancer therapy. Nat Rev Clin Oncol 16, 356–371. 10.1038/s41571-019-0175-7.

64. Moon, H., Donahue, L.R., Choi, E., Scumpia, P.O., Lowry, W.E., Grenier, J.K., Zhu, J., and White, A.C. (2017). Melanocyte Stem Cell Activation and Translocation Initiate Cutaneous Melanoma in Response to UV Exposure. Cell Stem Cell 21, 665–678 e666. 10.1016/j.stem.2017.09.001.

65. Zaidi, M.R., Davis, S., Noonan, F.P., Graff-Cherry, C., Hawley, T.S., Walker, R.L., Feigenbaum, L., Fuchs, E., Lyakh, L., Young, H.A., et al. (2011). Interferon-gamma links ultraviolet radiation to melanomagenesis in mice. Nature 469, 548–553. 10.1038/nature09666.

66. Handoko, H.Y., Rodero, M.P., Boyle, G.M., Ferguson, B., Engwerda, C., Hill, G., Muller, H.K., Khosrotehrani, K., and Walker, G.J. (2013). UVB-induced melanocyte proliferation in neonatal mice driven by CCR2-independent recruitment of Ly6c(low)MHCII(hi) macrophages. J Invest Dermatol 133, 1803–1812. 10.1038/jid.2013.9.

67. Gamba, C.A., Swetter, S.M., Stefanick, M.L., Kubo, J., Desai, M., Spaunhurst, K.M., Sinha, A.A., Asgari, M.M., Sturgeon, S., and Tang, J.Y. (2013). Aspirin is associated with lower melanoma risk among postmenopausal Caucasian women: the Women’s Health Initiative. Cancer 119, 1562–1569. 10.1002/cncr.27817.

68. Ogbechie-Godec, O.A., and Elbuluk, N. (2017). Melasma: an Up-to-Date Comprehensive Review. Dermatol Ther (Heidelb) 7, 305–318. 10.1007/s13555-017-0194-1.

69. Davis, E.C., and Callender, V.D. (2010). Postinflammatory hyperpigmentation: a review of the epidemiology, clinical features, and treatment options in skin of color. J Clin Aesthet Dermatol 3, 20–31.

70. Ding, Y., Xu, Z., Xiang, L.F., and Zhang, C. (2023). Unveiling the mystery of Riehl’s melanosis: An update from pathogenesis, diagnosis to treatment. Pigment Cell Melanoma Res 36, 455–467. 10.1111/pcmr.13108.

71. Kaufman, B.P., Aman, T., and Alexis, A.F. (2018). Postinflammatory Hyperpigmentation: Epidemiology, Clinical Presentation, Pathogenesis and Treatment. Am J Clin Dermatol 19, 489–503. 10.1007/s40257-017-0333-6.

72. Ebihara, T., and Nakayama, H. (1997). Pigmented contact dermatitis. Clin Dermatol 15, 593–599. 10.1016/s0738-081x(97)00072-2.

73. Taylor, A., Pawaskar, M., Taylor, S.L., Balkrishnan, R., and Feldman, S.R. (2008). Prevalence of pigmentary disorders and their impact on quality of life: a prospective cohort study. J Cosmet Dermatol 7, 164–168. 10.1111/j.1473-2165.2008.00384.x.

74. Ghasemiyeh, P., Fazlinejad, R., Kiafar, M.R., Rasekh, S., Mokhtarzadegan, M., and Mohammadi-Samani, S. (2024). Different therapeutic approaches in melasma: advances and limitations. Front Pharmacol 15, 1337282. 10.3389/fphar.2024.1337282.

75. Li, J.L., Goh, C.C., Keeble, J.L., Qin, J.S., Roediger, B., Jain, R., Wang, Y., Chew, W.K., Weninger, W., and Ng, L.G. (2012). Intravital multiphoton imaging of immune responses in the mouse ear skin. Nat Protoc 7, 221–234. 10.1038/nprot.2011.438.

76. Stuart, T., Butler, A., Hoffman, P., Hafemeister, C., Papalexi, E., Mauck, W.M., Hao, Y.H., Stoeckius, M., Smibert, P., and Satija, R. (2019). Comprehensive Integration of Single-Cell Data. Cell 177, 1888-+. 10.1016/j.cell.2019.05.031.

77. Stoeckius, M., Zheng, S., Houck-Loomis, B., Hao, S., Yeung, B.Z., Mauck, W.M., 3rd, Smibert, P., and Satija, R. (2018). Cell Hashing with barcoded antibodies enables multiplexing and doublet detection for single cell genomics. Genome Biol 19, 224. 10.1186/s13059-018-1603-1.

78. Aran, D., Looney, A.P., Liu, L., Wu, E., Fong, V., Hsu, A., Chak, S., Naikawadi, R.P., Wolters, P.J., Abate, A.R., et al. (2019). Reference-based analysis of lung single-cell sequencing reveals a transitional profibrotic macrophage. Nat Immunol 20, 163–172. 10.1038/s41590-018-0276-y.

79. Street, K., Risso, D., Fletcher, R.B., Das, D., Ngai, J., Yosef, N., Purdom, E., and Dudoit, S. (2018). Slingshot: cell lineage and pseudotime inference for single-cell transcriptomics. BMC Genomics 19, 477. 10.1186/s12864-018-4772-0.

